# Regulatory approved monoclonal antibodies contain framework mutations predicted from human antibody repertoires

**DOI:** 10.1101/2021.06.22.449488

**Authors:** Brian M. Petersen, Sophia A. Ulmer, Emily R. Rhodes, Matias F Gutierrez Gonzalez, Brandon J Dekosky, Kayla G Sprenger, Timothy A. Whitehead

## Abstract

Monoclonal antibodies (mAbs) are an important class of therapeutics used to treat cancer, inflammation, and infectious diseases. Identifying highly developable mAb sequences *in silico* could greatly reduce the time and cost required for therapeutic mAb development. Here, we present position-specific scoring matrices (PSSMs) for antibody framework mutations developed using natural human antibody repertoire sequences. Our analysis shows that natural human antibody repertoire-based PSSMs are consistent across individuals and demonstrate high correlations between related germlines. We show that mutations in existing therapeutic antibodies can be accurately predicted solely from natural human antibody sequence data. mAbs developed using humanized mice had more human-like FR mutations than mAbs originally developed by hybridoma technology. A quantitative assessment of entire framework regions of therapeutic antibodies revealed that there may be potential for improving the properties of existing therapeutic antibodies by incorporating additional mutations of high frequency in natural human antibody repertoires. In addition, high frequency mutations in natural human antibody repertoires were predicted *in silico* to reduce immunogenicity in therapeutic mAbs due to the removal of T cell epitopes. Several therapeutic mAbs were identified to have common, universally high-scoring framework mutations, and molecular dynamics simulations revealed the mechanistic basis for the evolutionary selection of these mutations. Our results suggest that natural human antibody repertoires may be useful as predictive tools to guide mAb development in the future.

## Introduction

Monoclonal antibodies (mAbs) are now ubiquitous as therapeutics, with over $100 billion in sales worldwide in 2020 (1) and applications ranging from oncology (2) and inflammation (3) to infectious diseases (4). mAbs are engineered not only to have potent and specific binding to a given target but also to have favorable drug properties, including *in vivo* stability, manufacturability, immunogenicity, solubility, and polyspecificity (5). Identifying highly developable mAb sequences *in silico* could greatly reduce the time and costs of therapeutic mAb development.

Natural antibody sequences sourced from human antibody repertoires could inform our ability to engineer therapeutic mAbs by ‘borrowing’ consensus mutations (6,7). This premise rests on the successful use of sequence conservation in protein engineering for improving the functional properties of enzymes (8–10), nanobodies (11), and membrane proteins (12). Antibodies in particular contain great potential for sequence optimization because every human body contains an estimated 10^11^ B cells with highly diverse antibody sequences (13), providing a rich space from which to glean important insights that could be used to guide future engineering efforts.

Using sequence conservation for improving antibody properties was first explored by Steipe et al. (14), who used known antibody sequences from the Kabat Database (15) to identify consensus positions within natural human antibody repertoires. Mutation to the amino acids at these consensus positions resulted in improved thermodynamic stability for the majority of the antibody sequences tested. However, the power of any sequence-based method relies on the size of the database. It is now possible to sequence tens of millions of antibodies from a single individual. Studies evaluating such human antibody repertoires have focused on cataloging the immune response to vaccination or infection (16–18). Recently, the Great Repertoire Project conducted the most extensive attempt to sequence entire human antibody repertoires to date, acquiring a total of 364 million antibody sequences by sequencing full Leukopaks from ten healthy, HIV-negative adults (19).

We revisited the idea that sequence conservation predicts developable antibody sequences using this much more comprehensive database of human antibody sequences, applied towards the analysis of FDA-approved mAbs. Specifically, we sought to answer the following questions: (*i.*) are germline (GL) mutations that are highly prevalent in natural human antibody repertoires also found in FDA-approved mAbs, given the generally favorable developability properties of these mAbs?; and more broadly (*ii.*) can sequence information alone predict more developable from less developable mAbs? We restricted our analysis to the framework (FR) regions of the variable heavy (V_H_) domain as antibody FRs impact *in vivo* stability, solubility, and immunogenicity (6) while also contributing significantly less than complementarity determining regions (CDRs) for binding antigen. We also explored some of the dynamics of peptide-MHC-II interactions using computational binding predictions (20), as the MHC-II peptide epitope contained within antibodies and other protein drugs has been recognized as an important component of clinical success (21,22). As a result, sequence information for FR regions can be applied to a broad array of antibodies with varying applications.

In this study, we present position-specific substitution profiles (PSSM for position-specific scoring matrix) for antibody FR mutations using the most complete dataset of human antibody repertoire sequences to date (19). We show that antibody repertoire-based PSSMs are consistent across subjects and produce high correlations between GLs with expected differences based on sequence similarity and familial relationships. Our analysis shows that mutations in existing therapeutic antibodies can be accurately predicted solely from repertoire sequence data. We then quantitatively assessed entire FRs of these therapeutic antibodies and compared them to their natural repertoire counterparts. These data suggest that there may be potential for improving existing therapeutic antibody properties through incorporation of additional mutations of high-frequency in natural repertoires. In addition, we found that high frequency repertoire mutations tended to reduce the affinity of germline-encoded peptides that bound to MHC-II epitopes. Several therapeutic mAbs have common, universally high scoring FR mutations, and simulations revealed a mechanistic basis for the favorable drug properties of some mutations. Overall, our results suggest that natural antibody repertoires are useful as predictive tools that will facilitate engineering mAbs by improving drug-like properties.

## Methods

### Selection of natural repertoire antibody sequences

In our analysis, we considered only the FR regions of immunoglobulin G (IgG) V_H_ antibody segments. IgM sequences are the other common isotype included in the database, but were ignored because they typically have low levels of mutation due to their role in the early stages of the immune response (23). D and J segments were excluded since these segments are highly variable, which would inhibit our ability to achieve significant coverage of all possible mutations at these positions.

Sequences for repertoire IgG variable heavy segments from a common previously-inferred GL were extracted from the Great Repertoire Project database (19) (https://github.com/briney/grp_paper) (data accessed November 2020) for each of the 25 GL genes analyzed. A minimum of 100,000 sequences were analyzed per GL. The cutoff value of 100,000 was deemed necessary for the creation of reliable scoring matrices, and was determined by a random subsampling analysis where score differences were compared across multiple sampling depths (**Figure S1**). It should be noted that the resulting compiled sequences span several different alleles within each GL gene, though these differences are later reconciled by adjusting the scoring matrices to remove this bias, as described in detail in a subsequent section. All GLs excluded from this analysis are in **Table S1**. GLs were excluded either due to low sequence counts or if no FDA-approved mAb precursors were represented in the panel.

**Table 1.** Statistical significance for framework mutations for FDA-approved mAbs.

| **Germline Gene** | **FDA-approved Antibodies** | **Number of V<sub>H</sub> Framework Mutations** | **p-value** | **Number of Repertoire Sequences Analyzed** |
| --- | --- | --- | --- | --- |
| VH1-2 | Pembrolizumab | 16 | 4.8E-06 | 300,000 |
| VH1-3 | Vedolizumab | 9 | 3.6E-02 | 129,913 |
| VH1-46 | Benralizumab<br>Ravulizumab<br>Burosumab | 24 | 9.3E-08 | 451,920 |
| VH1-69 | Risankizumab<br>Ixekizumab<br>Galcanezumab | 23 | 4.1E-06 | 720,993 |
| VH2-5 | Palivizumab | 7 | 4.4E-03 | 141,372 |
| VH3-7 | Secukinumab<br>Fremanezumab<br>Durvalumab | 10 | 5.8E-05 | 2,003,693 |
| VH3-9 | Adalimumab<br>Ofatumumab<br>Sarilumab | 9 | 1.0E-06 | 419,523 |
| VH3-23 | Pertuzumab<br>Denosumab<br>Daratumumab<br>Avelumab<br>Dupilumab<br>Emicizumab<br>Lanadelumab<br>Atezolizumab | 38 | 3.1E-07 | 300,000 |
| VH3-30 | Ipilimumab<br>Erenumab | 5 | 1.9E-04 | 1,058,653 |
| VH3-33 | Canakinumab<br>Nivolumab | 5 | 1.3E-02 | 710,048 |
| VH3-48 | Certolizumab pegol | 11 | 1.4E-03 | 300,000 |
| VH3-66 | Omalizumab<br>Trastuzumab<br>Eptinezumab | 27 | 2.6E-08 | 356,673 |
| VH3-74 | Efalizumab<br>Elotuzumab<br>Atezolizumab | 26 | 4.9E-05 | 1,355,540 |
| VH4-4 | Alemtuzumab | 8 | 2.1E-04 | 1,003,945 |
| VH4-30-4 | Tocilizumab<br>Necitumumab | 11 | 2.0E-06 | 160,757 |
| VH5-51 | Ustekinumab<br>Guselkumab | 7 | 2.2E-05 | 702,327 |

### Selection of FDA-approved antibody sequences

FDA-approved antibodies included in the analysis were selected according to three criteria: *(i.)* the sequences had to be human/humanized antibodies (name ending in -umab), *(ii.)* the sequences had to be available in the DrugBank database (24) at the time of sequence collection (accessed October 28th 2020), and *(iii.)* the Great Repertoire Project database (19) had to contain more than 100,000 sequences from the same inferred GL gene. It is critical that the analyzed antibody is human/humanized since this allows for the antibody to be matched to human GL genes.

In addition, to determine whether we can effectively identify developability issues in antibodies, we selected sequences for a second panel of engineered antibodies that were determined to have biophysical properties that fall outside of historically accepted limits. We used data from Jain et al. (5), which identified 12 biophysical properties that were indicative of an antibody’s likelihood of progressing through all three stages of FDA approval. From this dataset, we selected all antibodies that were found to have two or more biophysical properties that fell outside of the described acceptable ranges. In addition, the antibodies had to be human/humanized and the Great Repertoire Project database had to contain at least 100,000 sequences in the repertoire database from the same inferred GL gene. IMGT BlastSearch v1.2.0 (25) was used to infer the GL gene for the V_H_ sequence of each of the analyzed FDA-approved antibodies, and the hit with the lowest E value was assigned as the GL V_H_ gene. As described below, position-specific scoring matrices (PSSMs) for each of the 25 GL genes were then constructed (**Table S1**).

### Generation of position-specific scoring matrices for human V_H_ genes

A multiple sequence alignment was performed on each set of sequences derived from a common GL gene using a lightweight version of the MAFFT algorithm (26) designed for large numbers of similar sequences and using the FFT-NS-2 progressive alignment method in low-memory mode with the gap opening penalty set to max (5.0). All other alignment parameters were set to default. Each alignment file was collapsed into a single table of mutational counts by tallying the number of observations of each amino acid at each position in the sequence. This count output was modified to remove instances of insertions, indicated by positions that contained under 10% occupancy (i.e. more than 90% gaps). In addition, individual counts of wild-type (WT) and allelic variants were removed (**Table S2**). Amino acid substitutions with zero counts were replaced with a pseudocount of one to circumvent an undefined score (acting as a lower bound for scores). Then, the tables were manually aligned to IMGT human GL V_H_ FR reference sequences (25) to associate each column of the count table with a real position in the FR sequence.

**Table 2.** Differential stabilities for individual framework point mutants.

| V<sub>H</sub> Gene | Mutation (paper) | Mutation (IMGT) | Minimum nt | Score | Property | Reference |
| --- | --- | --- | --- | --- | --- | --- |
| V<sub>H</sub>3-23 | L20P | L21P | 1 | 12.0 | Mean $\Delta T_m = -2.1^\circ\text{C}^\ddagger$ | (46) |
| | Q81H | Q90H | 1 | 16.7 | Mean $\Delta T_m = 0.0^\circ\text{C}^\ddagger$ | |
| | A84P | A96P | 1 | 15.6 | $\Delta T_{50} = 0.7^\circ\text{C}$ | (47) |
| V<sub>H</sub>1-69 | S16E | S17E | 1-3<sup>†</sup> | -4.1 | $\Delta T_{50} = 9^\circ\text{C}$ | (48) |
| | S16Q | S17Q | 2-3<sup>†</sup> | -13 | $\Delta T_{50} = 5^\circ\text{C}$ | |
| | M48G | M53G | 2 | 0.4 | $\Delta T_{50} = 3^\circ\text{C}$ | |
| | M48I | M53I | 1 | 12.2 | $\Delta T_{50} = 2^\circ\text{C}$ | |
| | V67I | V76I | 1 | 15.1 | $\Delta T_{50} = 4^\circ\text{C}$ | |
| | V67L | V76L | 1 | 19.4 | $\Delta T_{50} = 7^\circ\text{C}$ | |
| V<sub>H</sub>6-1 | S16G | S16G | 2 | -0.9 | 180%<sup>a</sup>, 230%<sup>b</sup> yield* | (49) |
| | | | | | 6.2<sup>a</sup>, 7.3<sup>b</sup> kJ/mol $\Delta\Delta G$ | |
|  | V72D | V69D | 2 | 12.3 | 320%<sup>a</sup>, 180%<sup>b</sup> yield |  |
| | | | | | 0.1<sup>a</sup>, 2.2<sup>b</sup> kJ/mol $\Delta\Delta G$ | |
|  | S76G | S74G | 1 | 28.7 | 210%<sup>a</sup>, 150%<sup>b</sup> yield |  |
| | | | | | 3.7<sup>a</sup>, 3.5<sup>b</sup> kJ/mol $\Delta\Delta G$ | |
|  | S90Y | S88Y | 1 | 6.2 | 180%<sup>a</sup>, 230%<sup>b</sup> yield |  |
| | | | | | -0.1<sup>a</sup>, 1.4<sup>b</sup> kJ/mol $\Delta\Delta G$ | |

<sup>‡</sup>Mean $\Delta T_m$ was calculated over all genotypes presented in Table 3 in ref (28).
<sup>†</sup>Allele exists at this position.
\*Yield is calculated as a relative percentage to the original antibody sequence.
<sup>a</sup>:Antibody 2C2
<sup>b</sup>:Antibody 6B3

The counts for all amino acid substitutions within the FR regions were log transformed into a score via the following equation:

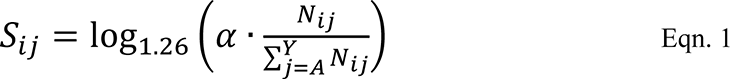

Here *S_ij_* is the score of amino acid substitution *j* at position *i* and *⍺* is a V_H_-specific constant (Range: 1021-2541, s.d.: 330) that centers the mean of the score distribution for V_H_-specific FR mutations at zero. For comparisons across GL families, this α normalization term results in a max difference of 3.9 between GL with a difference of <0.9 at 1 s.d. *N_ij_* is the count of amino acid substitution *j* at position *i*, the sum of which, over all amino acids from alanine (*A*) to tyrosine (*Y*), is equal to the total number of sequences without gaps in the multiple sequence alignment at that position. We used a log base of 1.26 to adhere to previous convention used by Dayhoff et al. for log-odds matrices (27). PSSMs for the 25 analyzed GLs can be found in **Data S1**. This same protocol was then applied to the FDA-approved antibody sequences, which were first aligned to their corresponding GL sequences and then nonsynonymous FR mutations and their respective scores were recorded.

### Generation of framework scores for individual antibody sequences

We then derived an overall FR region score that we term an “FR score”, a metric which can be used to compare sequences of antibodies with different numbers of mutations across different GLs. We assumed that each FR mutation has an additive effect and is independent of one another (no epistatic interactions). With this assumption, the FR score is then defined as:

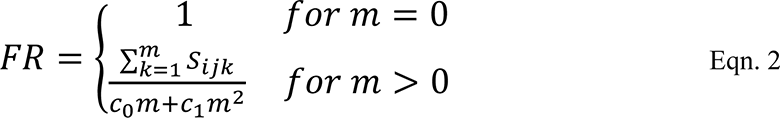

where *S_ijk_* is the score of the *k*^th^ sequential FR mutation of amino acid substitution *j* at position *i* as determined by Eqn. 1. The sum of *m* (the number of FR mutations) mutation scores are normalized by constants *c_0_* and *c_1_* which are specific to that GL gene (**Table S3**), derived from a weighted least squares regression. The denominator of the FR score is a normalization constant which gives an estimate of a typical score of a randomly sampled Ab from the repertoire with the same number of FR mutations (**Figure S5**). The FR score as derived gives an estimation of how different an antibody’s framework mutations are as compared to natural repertoire antibodies. As such, if an antibody has framework mutations with scores similar to what we would expect of a natural antibody with an equivalent number of mutations, this ratio becomes one. Thus, we assign all antibodies with no framework mutations an FR score of one.

### Prediction and analysis of MHC-II peptide epitope affinity

MHC-II peptide epitope affinity prediction was carried out for a set of 38 representative HLA-DRB1 alleles (20) using netMHCIIpan 4.0 (28). Predictions were performed using 15-mer peptide scans for each peptide containing mutations with an associated PSSM, and the netMHCIIpan flag -BA was selected to obtain predicted MHC-II binding affinities. Predictions were also carried out for germline (GL) peptides using the same settings, for the 38 representative HLA-DRB1 alleles. Next, netMHCIIpan predictions for FDA-approved mAb peptides were matched to GL peptides and their own binding predictions. To track affinity changes from high affinity GL peptides, peptides matching a germline peptide with peptide:MHC-II K_D_ < 1,000 nM were assigned to three different bins based on the ratio of (mutated peptide K_D_ / GL peptide K_D_). This was done for KDs predicted for the same allele. Three bins were created: K_D_ fold-change < 0.5, 0.5 ≤ K_D_ fold-change ≤ 2, and K_D_ fold-change > 2. We also considered the total set of peptides matching any high affinity GL peptides (K_D_ < 1,000 nM), and the total set of peptides matching any low affinity GL peptide (K_D_ > 1,000 nM). Next, PSSM values were averaged by DRB1 allele and K_D_ group using standard mean. This was done to reduce redundancy, since the same peptide can have different affinities for different HLA-DRB1 alleles and thus be in different K_D_ groups at the same time. As a result, average PSSMs per DRB1 and K_D_ group take into account peptide contributions based on differential DRB1 binding. Next, PSSM distributions were compared between K_D_ bins using a Welch Two Sample t-test, and p-values were adjusted for multiple comparisons using the Benjamini-Hochberg method. All data processing steps were carried out in R.

### Molecular dynamics simulations

Molecular dynamics (MD) simulations were carried out using the GROMACS 2021 (29,30) MD engine and the TIP3P (31) and AMBER99SB-ILDN (32) force fields to explicitly model water and the antibodies, respectively. Simulations were initiated from crystal structures of the following antibodies: Atezolizumab (PDB code 5X8L) (33), Daratumumab (PDB code 7DUN) (34), and Omalizumab (PDB code 4X7S) (35), with all antigens/ligands removed beforehand. Additional simulations were performed with a single germline reversion mutation incorporated into each mAb sequence (A54S for Atezolizumab and Omalizumab, and F103Y for Daratumumab), using Visual Molecular Dynamics (VMD) (36). Counterions in the form of Na^+^ or Cl^-^, also described by the AMBER99SB-ILDN force field, were added to the systems as needed to neutralize any net charge. Each system contained between 67,000 and 94,000 atoms.

The initial coordinates for each system were minimized for a maximum of 5,000 steps using a steepest descent energy minimization. This was followed by 0.5 ns each of NVT and then NPT equilibration simulations performed at 310 K using the Bussi−Donadio−Parrinello thermostat (37), and at 1.0 bar using the Berendsen barostat (38) (same temperature and thermostat) for the NPT simulations. The time constant for coupling in both the NVT and NPT simulations was 0.1 ps. NPT production simulations were then performed at the same temperature and pressure and using the same thermostat and Parrinello-Rahman barostat (39). Particle mesh Ewald (PME) summations (40) were used to calculate long-range electrostatic interactions with a cutoff of 1.0 nm, and Lennard Jones interactions were calculated over 1.0 nm and shifted beyond this distance. Neighbor lists were updated every 10 steps with a cutoff of 1.0 nm and all simulations utilized full periodic boundary conditions. Production simulations were carried out for 0.3 *μs* each, for a total of 1.8 *μs* across the six simulations.

Standard GROMACS tools were used to compute the root mean squared deviation (RMSD) of the antibodies from their respective energy-minimized structures as a function of simulation time. For all RMSD plots, data was plotted every 10^th^ frame. We also computed the root mean squared fluctuation (RMSF) of the heavy chains of the antibodies over the last half (150 ns) of the simulations, during which time all six simulations were deemed converged based on RMSD. All simulation images were rendered in VMD.

### Statistical calculations

Unless otherwise noted, all p-values were calculated using Welch’s t-test. Correlations were analyzed using a Pearson product-moment correlation coefficient. Hierarchical clustering for GL comparisons was performed using SciPy in Python (www.scipy.org) using the unweighted pair group method with arithmetic mean (UPGMA).

## Results

### 3.1 Generation of unbiased framework PSSMs from antibody repertoires

We created an efficient method to score the set of individual FR mutations in an FDA-approved mAb, which is shown schematically in **Figure 1** and is carried out as follows. First, we identified the set of human or humanized FDA-approved mAbs for which sequences were available in the DrugBank database (24). From this list, we used IMGT BlastSearch (25) to infer a GL V_H_ gene for each mAb (**Table S1**). We restricted our analysis to the FR positions of V_H_ only (IMGT Numbering FR1-26; FR39-55; FR66-104), as the Great Repertoire Project reported only heavy chain sequences. Additionally, we considered only IgG sequences, as IgM sequences typically have low levels of mutation due to their role in the early stages of the immune response (23).

**Figure 1.**
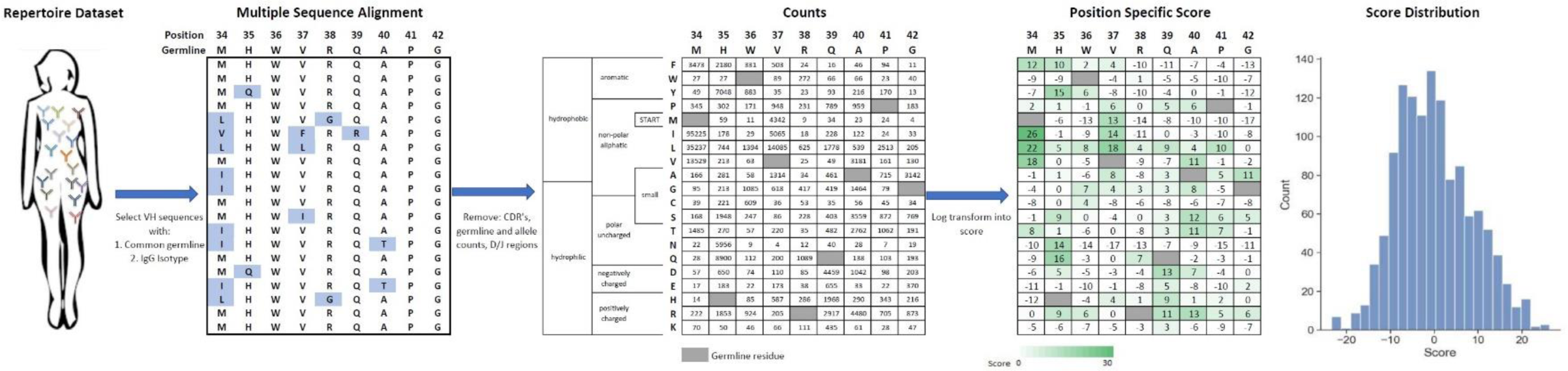
Generation of V_H_ germline position specific scoring matrices. IgG sequences from a specific V_H_ germline are extracted from the repertoire dataset and aligned. Mutations from germline in V_H_ framework regions are tabulated and then log-transformed into a position-specific score. This score function centers the population of scores at zero with more frequent mutations indicated by a higher score. Data shown here is for the VH1-2 germline.

We aligned a set of relevant IgG sequences by multiple sequence alignment (26) in order to generate a FR position-specific scoring matrix (PSSM) for each GL V_H_ gene considered (see **Methods)**. For each FR position, we counted the number of sequences observed to have a mutation at this position and log-transformed this frequency to a score term. This scoring term has one adjustable parameter per GL V_H_ gene such that the mean score for a randomly selected mutation would be zero. According to how we defined this scoring term (see **Methods**), higher scores represent more commonly observed amino acid mutations and the gross majority of scores range between -10 and 10.

One complication arising from statistical analysis of antibody mutations is that humans possess significant allelic diversity in V_H_ genes, even in FR regions. For example, consider V_H_3-9 which has a T99M FR allelic variation (**Table S2**). In generating a PSSM at position 99, one could identify the appropriate allele for each patient (either T or M) and then generate separate PSSMs for each allelic variation. Alternatively, we chose the simpler approach of removing all allelic variants from our PSSM under the reasonable assumption that such alleles would be relatively well tolerated in mature mAbs and should not result in large changes in antibody properties. All identified alleles were present in the IMGT database with one exception: we noticed that one VH1-69 mutation (S17A) occurred at a frequency approaching 90%. We reasoned that this mutation most likely represents an allele not currently represented in the IMGT database and so excluded that S17A mutation from further analysis.

The actual number of IgG GL V_H_-specific sequences in the Great Repertoire Project database varies tremendously, from around 2,500 sequences (VH1-45) to more than 3.9 million sequences (V_H_3-23). Given this range, we asked what number of sequences would be sufficient for accurate recapitulation of GL PSSMs. We chose to perform this analysis on VH5-51 by subsampling between 1,000 and 300,000 sequences. For each mutation, we computed the absolute difference between the score derived from the full dataset (∼700,000 sequences) and the score from the subsample. Increasing the number of sequences from 1,000 to 25,000 led to increasing agreement with the full dataset, with essentially no difference observed in the median displacement at or above 25,000 sequences for the large majority of scores (i.e., scores greater than -15) (**Figure S1**). However, minor differences in displacement at 2σ (95%) were observed between 25,000 and 100,000 sequences. Thus, we conservatively restricted our analysis of GL V_H_ segments to those with 100,000 or more IgG sequences present in the Great Repertoire Project database. The GLs analyzed and resulting sequence counts are denoted in **Table S1**.

We then asked whether mutation frequencies would be highly correlated between individuals and if the site-specific preferences would be repeatable. To test this, we generated patient-specific V_H_3-23 PSSMs for all 8 individuals for which data was available which contained the same V_H_3-23*01 allele. The V_H_3-23 GL was selected to ensure that a sufficient number of sequences was available for generating these PSSMs. We observed a high degree of correlation between patient scores with correlation coefficients between 0.86 and 0.91 for all pairwise comparisons (**Figure 2**). We then repeated our analysis considering only amino acid mutations encoded by 1-nucleotide (nt) substitution, as these points contain the highest sequence coverage in our dataset and, therefore, are presumably most accurate. The results were essentially unchanged, with pairwise correlation coefficients between individuals ranging between 0.85 and 0.92 (**Figure S2A**). These correlation coefficients are only slightly below the basal noise level (0.94) calculated by computing the correlation coefficient between two independent random samples for each of two patients (**Figure S2B**). These results show that GL V_H_-specific FR mutations are largely repeatable between individuals. This finding is not new, as Sheng et al. showed in their analysis that affinity maturation somatic hypermutation (SHM) seems to produce highly consistent substitutions for antibody V segments across different donors, even in the absence of functional selection (16). However, our independent analysis on a unique dataset corroborates their results, which together suggest underlying biophysical mechanisms are responsible for the reproducibility of the selection of certain amino acid mutations at different FR positions. These results strongly suggest that the process of affinity maturation results in replicable frequency distributions of FR mutations at the amino acid level.

**Figure 2.**
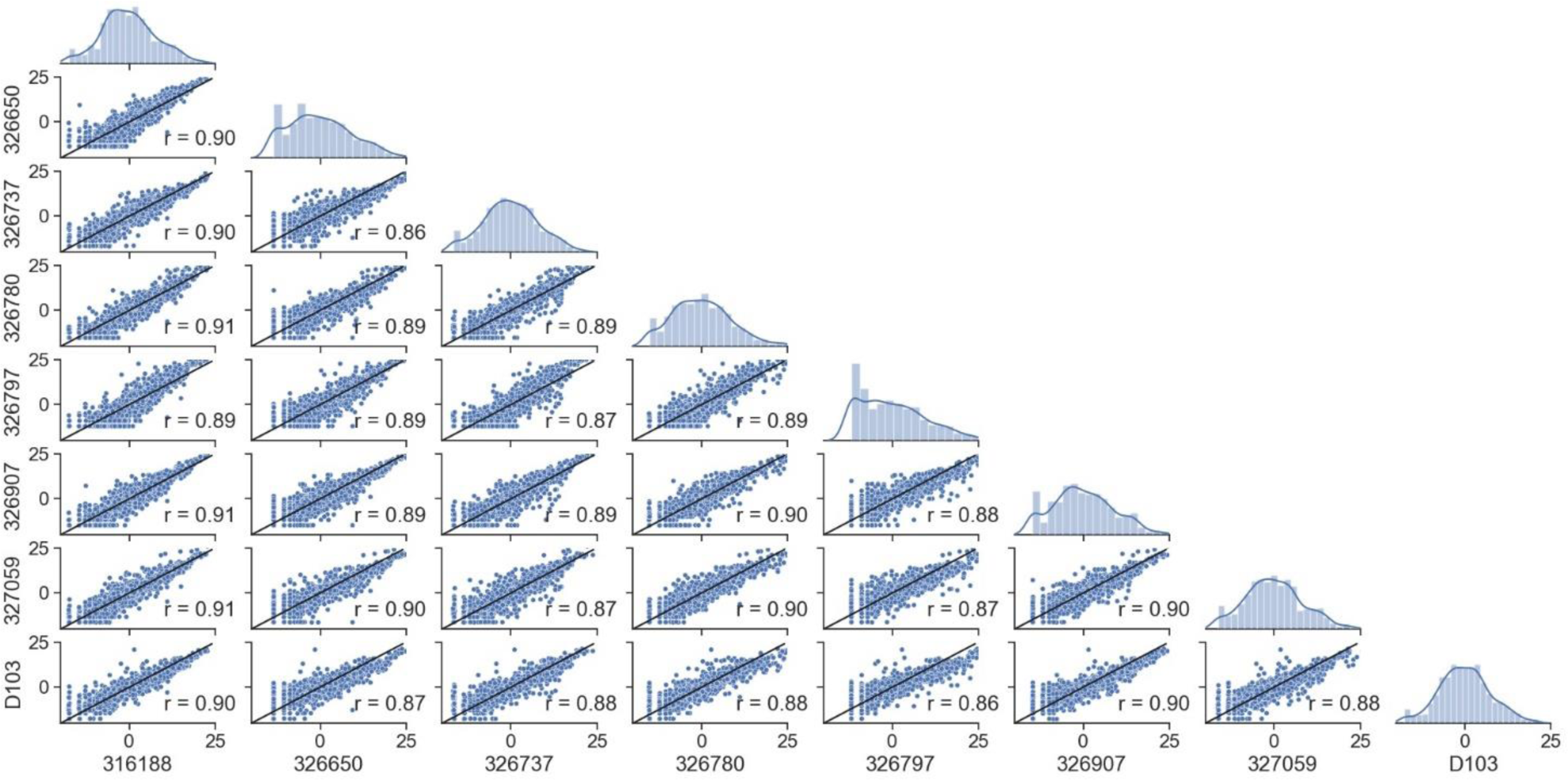
Framework scores are repeatable between individuals. Distributions and correlations of patient-specific PSSM scores for the V_H_3-23*01 germline. Alphanumeric characters on the y- and x-axes represent de-identified subjects.

Such replicable frequencies can arise if a given FR mutation is selected during affinity maturation for a general, antigen-independent reason. For example, a given FR mutation may improve the surface density of B cell receptors, or could remove an aggregation-prone patch on the surface of the protein. An alternative but non-exclusive explanation is as follows. Sequences from natural antibody repertoires tend to be enriched in mutations which can be generated by a single nucleotide substitution, while mutations that would require di- or tri-nucleotide substitutions tend to occur less frequently. This is because activation-induced cytidine deaminase (AID) primarily generates single nucleotide point mutations in B cell receptor sequences during affinity maturation (41), and as this mutational frequency is ∼10^-3^ per base per generation (42), it is far less probable that the same codon will be targeted twice or thrice than once. Consistent with this notion, analysis of our PSSMs shows that FR mutation scores that are one nucleotide away from the GL sequence are significantly higher (mean score of 8.50) than those requiring two or three substitutions (mean scores of -2.28 and -8.08, respectively) with two exceptions (VH1-24, VH1-46) (**Figure S3**).

### 3.2 FDA-approved mAbs contain framework mutations that can be predicted from natural antibody repertoires

We asked whether FR mutations in FDA-approved mAbs could be predicted from native antibody repertoires. 39 FDA-approved human/humanized mAbs were found to have sufficient depth of coverage (i.e., > 100,000 sequences) in their GL V_H_ family in the Great Repertoire Project database, and were thus subjected to further analysis (**Table 1**). While the precise developmental pathways for all of these FDA-approved mAbs are not publicly available, it is known that most of the mAbs were discovered using either hybridoma technology, transgenic mice, or phage display.

On average, the 39 FDA-approved mAbs have a mean of six V_H_ FR mutations, and a range from zero to 16 FR mutations. The degree to which these mutations might align with (and thus may be predicted by) those found in natural antibody repertoires is difficult to know *a priori*. On the one hand, the use of phage display or transgenic mice could result in a sufficiently different selection environment than what exists during affinity maturation in human germinal centers, leading to an altogether different set of FR mutations than are evolved in natural antibody repertoires. On the other hand, the clinical approval process eliminates any antibodies that may have been selected ineffectively for drug-like properties and failed in manufacturing or pre-clinical and clinical trials. The affinity maturation process may indirectly select for some of the same qualities of antibodies that have gone through regulatory approval, including high expression yield, thermodynamic stability, low non-specific binding, little to no aggregation propensity, and low immunogenicity (43,44). In this latter case, FR mutations in FDA-approved mAbs could indeed be predicted by natural antibody repertoire abundance.

To answer this question, we used our GL V_H_-specific PSSMs to score each FR mutation contained in the FDA-approved mAbs. Strikingly, every one of the 16 GL V_H_ families that form the set of inferred precursors for these mAbs showed significantly higher (at 95% confidence level) FR mutation scores for FDA-approved mAbs relative to the set of all possible mutation scores (**Figure 3A-B**, **Figure S4, Table 1**). The statistical significance of this result ranged from a p-value of 4.1e-6 (VH1-69) to 2.6e-8 (V_H_3-66) (**Figure 3B**). However, the distributions in **Figure 3B** of the set of all possible mutation scores do not reflect the *number* of repertoire antibody FR mutations with given scores. Indeed, generating random samples of scores for 1,000 repertoire antibody sequences from each of these distributions shows that the majority of observed antibody mutation scores in human repertoires are above 10 (**Figure 3C**). Plotting the individual mutation scores for the FDA-approved mAbs on top of these new distributions, we see that the distributions match much more closely, with scores of the FDA-approved mAbs now slightly below those of the natural antibody repertoires (**Figure 3C**). The slightly lower scores of the FDA-approved mAbs largely result from the fact that in each case, there are at least two mutations below a score of zero, and for the entire dataset of FDA-approved FR mutations 9.8% had a score below zero. These results suggest that while repertoire datasets can predict mutations in mAb V_H_ FR regions with reasonable accuracy, as indicated by the large overlap in the distributions, there is also considerable room for engineering mAb sequences to mimic natural repertoire antibodies with increased fidelity.

**Figure 3.**
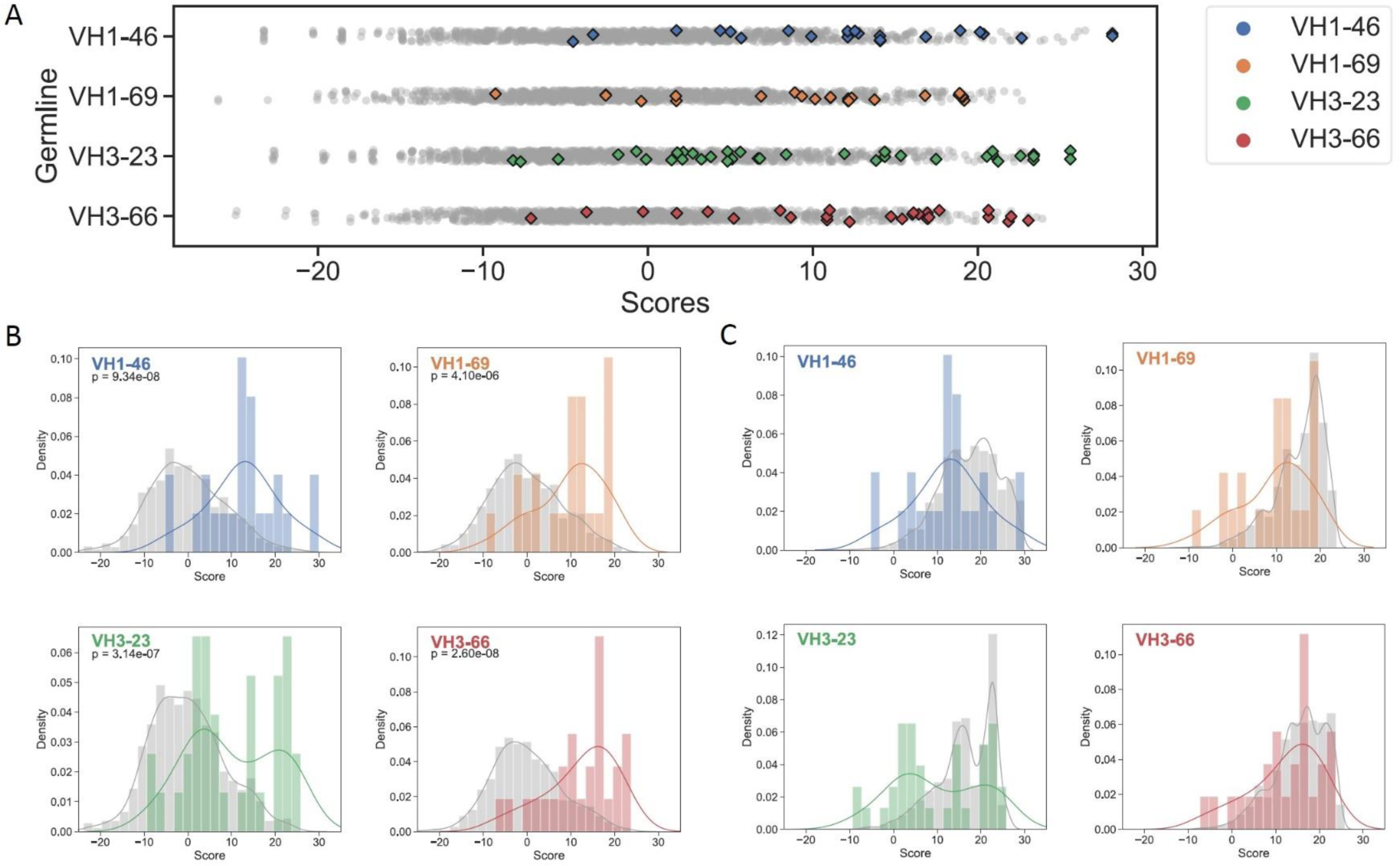
Regulatory approved monoclonal antibodies contain framework mutations predicted from human antibody repertoires. Repertoire-based PSSM scores (gray) compared to FDA-approved FR mutation scores (colored) for each of four germline genes (Blue: VH1-46, Orange: VH1-69, Green: V_H_3-23, Red: V_H_3-66). Individual scores (A) and distributions with kernel density estimates (B-C) are shown for the entire set of repertoire scores (B) and all FR mutation scores for a sample of 1,000 repertoire antibody sequences (C). P-values are calculated using single-tailed Welch’s t-test.

### 3.3 FDA-approved mAbs have lower FR scores than typical repertoire antibodies

Motivated by the previous results, we wondered how the cumulative effect of all FR mutations for a given FDA-approved mAb compares to that of natural repertoire antibodies. To assess this, we assumed, for simplicity, that individual mutations are independent, such that the cumulative effect of FR mutations is best represented as a sum of individual mutation scores. Due to the V_H_-specific centering parameter included in the score function, individual mutation scores cannot be directly compared across V_H_ families. To account for these differences, the score sum was normalized by a regression function which estimates the typical score sum of a randomly sampled repertoire antibody from the same V_H_ family with the same number of FR mutations (**Figure S5** and **Table S3**). The resulting “FR score” is defined such that a score of one represents an antibody that has FR mutations similar to that of a typical repertoire antibody framework sequence (see **Methods**). Antibody sequences with no FR mutations were assigned a pseudo-score of one since the calculated FR score is otherwise undefined.

We then compared FDA-approved mAbs to repertoire antibodies using FR scores. A simple random sample with replacement of 1,000 repertoire antibody sequences across all GL genes was generated and an FR score for each sampled antibody was computed (**Data S2**). As expected, FR scores for the repertoire antibodies center at a value of one (**Figure S6**). Next, FR scores were calculated for all FDA-approved mAbs used in the analysis above and the results are plotted along with the repertoire averages in **Figure 4A** as a function of the number of FR mutations. All individual mutation scores and FR scores for FDA-approved mAbs are tabulated in **Table S4**.

**Figure 4.**
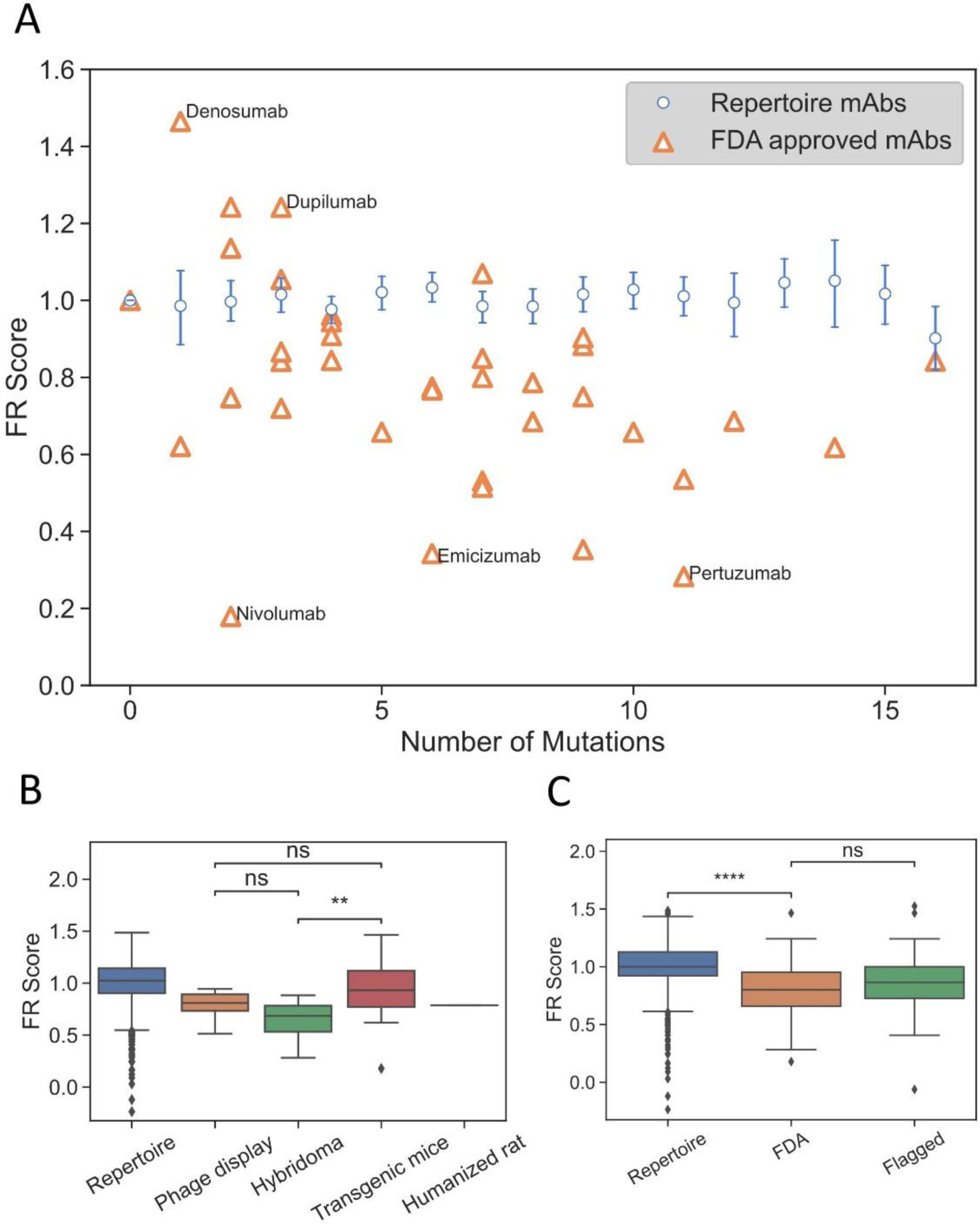
FR scores of FDA-approved and flagged mAbs are lower than typical human antibodies. (A) Comparison of FR scores for repertoire Abs and FDA-approved mAbs. Error bars on FR scores represent 95% confidence intervals. (B) FR scores of FDA-approved mAbs by development method. (C) FR scores of repertoire, FDA-approved, and flagged mAbs. (ns: p > 0.05, **: p < 1e-2, ****: p < 1e-4)

Thirty of the 39 FDA-approved mAbs have FR scores below the natural repertoire antibody average of one (p-value 2.7e-5). **Figure 4A** highlights the five mAbs with the highest and lowest FR scores. Denosumab and Dupilumab are the two highest scoring FDA-approved mAbs with FR scores of 1.46 and 1.24, respectively. Both mAbs have a small set of apolar/polar FR mutations (A55G, A25G, S40T, A55S), and interestingly, both mAbs share the same V_H_3-23 GL and were produced from transgenic mice. Nivolumab is the lowest scoring FDA-approved mAb with an FR score of 0.18 and only two mutations requiring more than 1-nt change from the GL sequence (S22D: score=4.7; A24K: score=2.3). This is unexpected since these scores are far below the average for 1-nt change mutations (average=8.50). Emicizumab and Pertuzumab, both from the V_H_3-23 GL and both developed using hybridoma technology, also have low FR scores of 0.34 and 0.28, respectively. However, the low scores of these mAbs are only somewhat explained by the fact that 14 mutations (out of 17 total) are more than 1-nt away from the GL codon. The solved complex of Pertuzumab and human epidermal growth factor receptor 2 (ErbB2 or HER2) (45) partially reveals the source of Pertuzumab’s low FR score. The complex is unusual in that FR positions surrounding CDRH2 are directly contacting ErbB2, including mutations for Y66I, A68N, D69Q, and N82R. Thus, rare FR mutations are necessary in this specific mAb for Erb2 recognition.

As shown above, the methods used to develop mAbs can influence the selective pressure and, therefore, the mutations that are incorporated into therapeutic mAbs. On average, we find that the number of FR mutations in FDA-approved mAbs discovered by hybridoma technology is higher than in FDA-approved mAbs discovered by transgenic mice or phage display (p-value ≥5.5e-7) (**Figure S7**). Even though FR scores are independent of number of FR mutations, we find that mAbs developed using hybridoma technology scored significantly lower than transgenic mice (p-value 8.3e-3), although there is not a statistically significant difference between mAbs developed via hybridoma versus phage display (p-value 0.10) (**Figure 4B**). Phage display and transgenic mice technologies produce mAb FRs with similar FR scores (p-value 0.20). Thus, for existing mAbs, those produced by transgenic mice are the most representative of the natural human antibody repertoire in terms of their FR mutations.

We next questioned whether FR scores could be used to diagnose mAb developability issues. Collecting an appropriate dataset for this analysis is difficult, as there are few databases of therapeutic antibodies with developability issues available in the publicly accessible literature. As a proxy, we chose to use the set of “flagged” antibodies originally described in Jain et al. (5). Jain et al. examined 12 different biophysical properties of a set of mAbs in various stages of clinical trials, with outliers in any of these properties considered “flags”. Two or more of these flags identified a mAb as a developability risk. We compiled all human/humanized mAb sequences from this analysis that contained at least two flagged properties and sufficient repertoire sequence data (39 in total). It should be noted that five of the flagged Abs that we analyzed have since received FDA approval and are therefore also contained within the FDA-approved dataset. FR scores were calculated for flagged mAbs and compared to our previously described panel of FDA-approved mAbs (**Figure 4C**, **Figure S8**, **Table S5**). Six of the flagged mAbs contained no FR mutations. As with the FDA-approved mAbs, we see that flagged mAbs also contain significantly lower FR scores than a population of randomly selected repertoire Abs (p-value 8.9e-3). However, there is no statistically significant difference between the distribution means (p-value 0.25) for regulatory-approved vs. flagged mAbs. Thus, our FR score is too coarse-grained to differentiate between antibody groups at such a late stage of development.

### 3.4 Biophysical basis of highly prevalent FR mutations

Why is the frequency of a given FR mutation largely repeatable between individuals? It could be the case that AID preferentially encodes given FR mutations due to sequence hotspots (41), with minimal selection of the amino acid sequence. Alternatively, there may be underlying biophysical bases for the selection of certain sequences. However, uncovering the rationale for every FR mutation is challenging as any individual mutation is pleiotropic, thus impacting many qualities *in vivo* including aggregation propensity, proteolytic stability, B cell receptor expression, antigen-binding affinity, and polyspecificity. Beyond these traits, it has also been hypothesized that engineering antibodies with FR regions derived from GL gene-preferred substitution patterns would result in lower immunogenicity as these antibodies would more closely reflect native antibody responses, which are readily recognized by the immune system (6,20). As an initial effort, we decided to focus on two properties which could reasonably explain the molecular basis behind these high scores: stability and MHC-II peptide epitope content. These biophysical properties can be addressed sufficiently using *in vitro* biochemical assays or *in silico*.

Other research groups have published the effects of FR mutations on stability and heterologous expression yields (**Table 2**) (46–49). Across these four studies, thermodynamic stability was measured (as ΔΔG, ΔT_50_, ΔT_m_) for 13 point mutants. Seven of the eight point mutants required a 1-nt substitution and had an individual mutation score above zero. These seven showed neutral or improved stability. Additionally, the two mutants with the highest mutation scores (V_H_1-69: V76L; V_H_6-1: S74G) also showed the highest thermodynamic stability (measured as ΔT_50_). Together, these results hint at a relationship between stability and repertoire abundance of a given FR mutation, although the small size of this dataset does not allow for sufficient statistical power to definitively draw a link between these factors.

FR mutations could also be selected to decrease the MHC-II epitope content of a given antibody sequence. Previous research has shown that unmutated antibody sequences have germline-encoded peptides that bind to and are presented by MHC-II proteins. These peptide epitopes are targeted for removal by SHM during B cell development *in vivo* (20). We sought to determine whether individual FR PSSM scores calculated here correlate with differential peptide:MHC-II binding affinities. To address this question, MHC-II binding affinities (KD) for all 15-mer peptides in a sliding window (i.e., fragments of 15 consecutive amino acids) sourced from the 39 FDA approved mAbs were calculated using a custom computational pipeline (see **Methods**). For each peptide containing FR mutations, mutation scores were binned by the K_D_ fold-change from the corresponding unmutated peptide’s K_D_. Interestingly, peptides with > 2 K_D_ fold-change from their unmutated germline peptides showed higher PSSM scores than mutations with < 0.5 K_D_ fold-change, and these data were statistically significant. This analysis suggests that prevalent FR mutations are more likely to decrease the affinity of mAb peptide fragments for the MHC-II groove than to increase the peptide:MHC-II affinity at a given mutation site; these data were consistent with prior reports of natural repertoire antibodies using paired heavy and light chain human sequence datasets (20) (**Figure 5**). As expected, mutations with moderate or no change in binding (0.5 ≤ K_D_ fold change ≤ 2) showed the highest PSSM scores across all groups because the majority of mutations do not affect the ∼5-8 heavy chain variable region MHC-II peptide epitopes for a given MHC-II gene. Strikingly, we found that the high-affinity MHC-II peptide epitopes showed higher PSSM scores than low-affinity MHC-II peptide epitopes, and the differences were statistically significant. Together, these data demonstrate that high-affinity MHC-II peptide epitopes inside the antibody heavy chain variable region are preferentially targeted for mutations, in both natural repertoires and in clinically approved mAbs (**Figure 5**).

**Figure 5.**
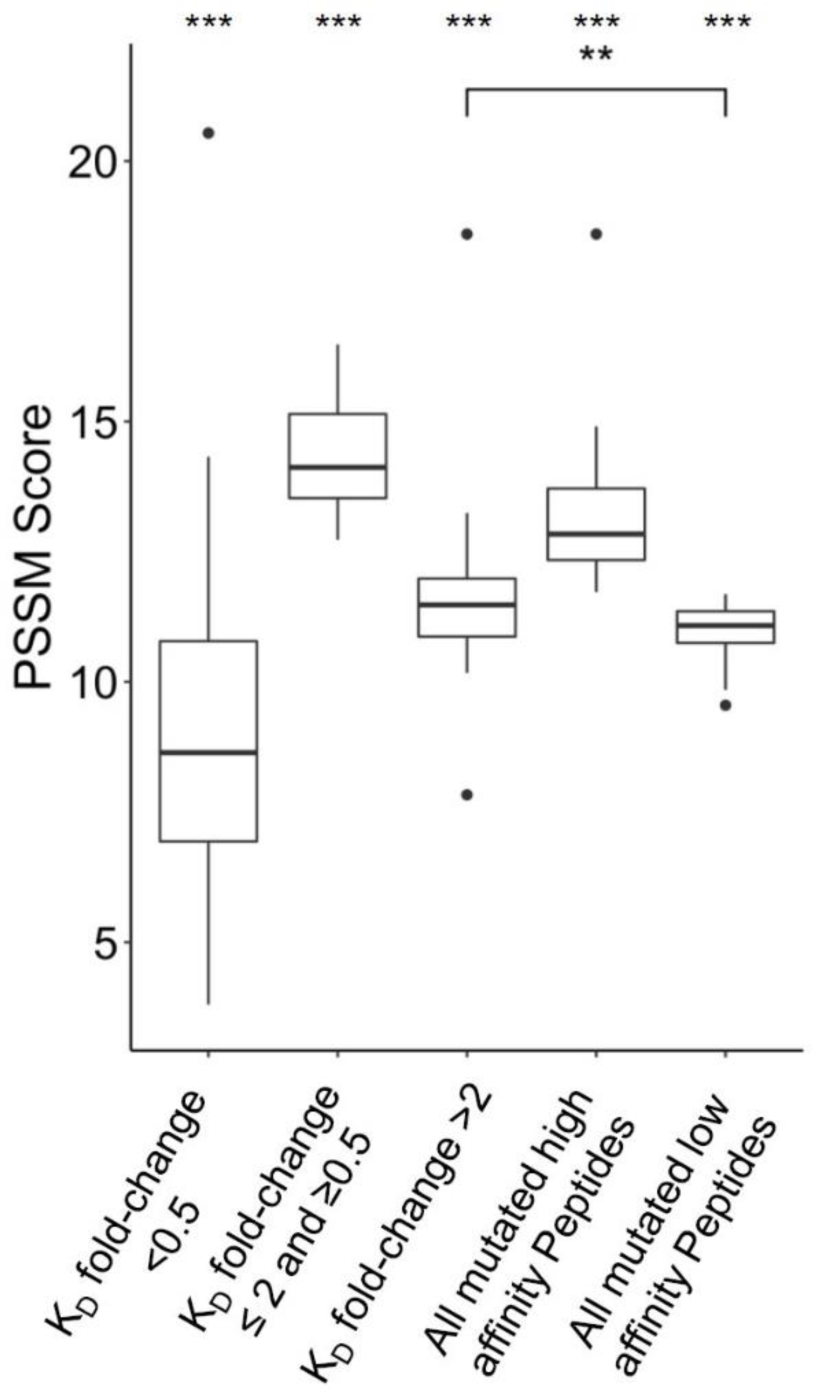
Position-Specific Scoring Matrix (PSSM) distributions for peptide mutations present in FDA-approved mAbs, binned by peptide:MHC-II K_D_ changes. Peptides were matched to V-gene germline peptides after peptide K_D_ prediction. For each group, the mean PSSM scores for each DRB1 allele with median and interquartile ranges are shown. Peptides mutated from high affinity germline peptides (K_D_ < 1,000 nM) were grouped based on K_D_ fold-change from germline. The complete set of high affinity and low affinity peptides (K_D_ < 1,000 nM) is also shown. Differences between groups was determined by a Welch Two Sample t-test, and p-values were adjusted for multiple comparisons. A total of 3,135 peptides were analyzed. ***: p-value < 0.001 for comparisons between all groups, except when noted. **: p-value < 0.01

### 3.5 Position-specific framework mutation scores are highly correlated between germline families

It has been established that affinity maturation produces antibodies that have GL-specific substitution patterns (16). What is the exact correlation of substitutions between GL V_H_ genes? To address this question, we calculated the Pearson correlation coefficient between each pair of GL PSSMs (**Figure 6**). Overall, correlation coefficients were quite high between GL PSSMs, ranging from 0.50 to 0.95. Hierarchical clustering was performed, revealing that correlations within subfamilies were higher than those between subfamilies. For example, all of the V_H_1 GL members clustered tightly together, as did those in V_H_3, and in V_H_4. The resulting clusters may, to an extent, be reflective of sequence similarity: GL members within a subfamily will have a higher pairwise amino acid identity than between subfamilies, which could skew the results. To account for this, we also calculated Pearson correlation coefficients only between GL PSSMs for which a common DNA codon is shared (**Figure S9**). As expected, higher correlations were observed between subfamilies than before, but the overall clustering within subfamilies was conserved. Thus, FR mutations are largely, though not wholly, repeatable between GL families and the correlation is stronger within subfamilies than between subfamilies.

**Figure 6.**
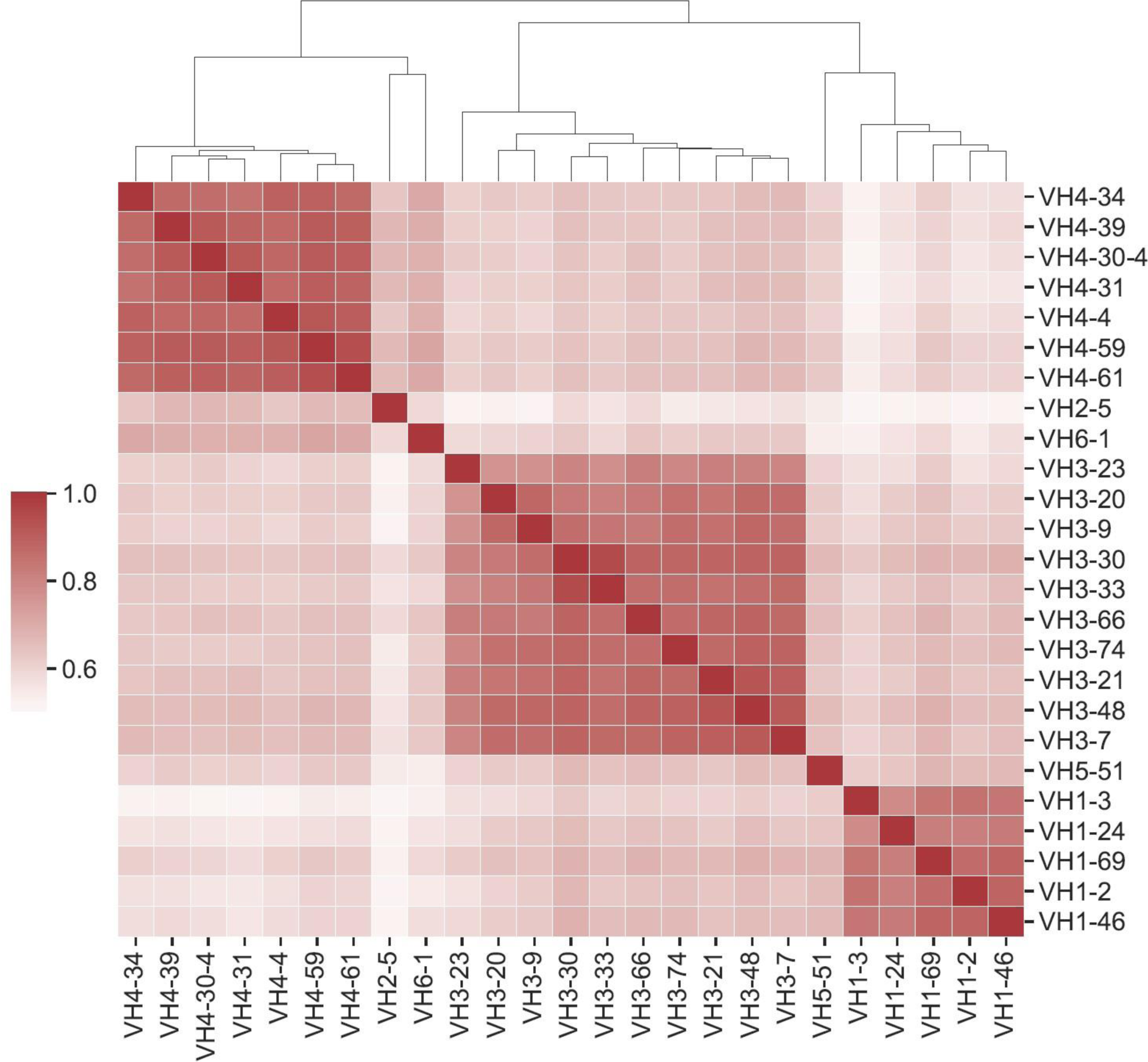
Position-specific framework mutation scores are highly correlated within V_H_ germline gene families. Pearson correlation coefficients were calculated for each pair of PSSM scores. Germline families are grouped by hierarchical clustering of Pearson correlation coefficients. Dendrogram indicates similarity between germline members.

### 3.6 Certain highly prevalent FR mutations are observed across many germline families

We next looked at ‘universal’ FR mutations, or highly abundant mutations that are seen across all subfamilies considered. 18 mutations at 14 FR positions were enriched across all GL members scanned in this work (**Table S6**). Overall, ‘universal’ mutations were largely chemically similar and/or polar substitutions at positions predominantly on the protein surface and distal to the CDRs. ‘Universal’ mutations also tended to be the WT residue for at least some of the GL families. For example, serine at FR85 is highly prevalent across GL families and is the WT residue for several V_H_ GL families (V_H_1-2, V_H_1-3, V_H_1-46, V_H_1-69, V_H_5-51).

We noticed a specific mutation (S54A) that shows up in several FDA-approved mAbs (Pertuzumab, Atezolizumab, Omalizumab, and Trastuzumab) with high scores across multiple GL V_H_s (V_H_3-23 and V_H_3-66 with scores of 21 and 17, respectively). **Figure 7A** shows the predicted scores for the mutation of residue 54 to five different amino acids (A, C, G, S, and T) across all 25 GLs studied. We observe that this specific mutation also shows up in multiple other GL V_H_s for which there is no corresponding FDA-approved mAb (V_H_3-20, V_H_3-21, V_H_3-48, V_H_3-74, and V_H_3-9, with scores ranging from 20-22). To understand the mechanistic basis for S54A selection, we analyzed the four FDA-approved mAbs listed above with this S54A mutation. To test for increased global stability of the mAbs, we utilized the online protein stability prediction algorithm PremPS (50) to determine the impact of reverting the alanine at position 54 in each of the four mAbs back to its identity of a serine in the respective germline sequences (**Figure 7B**). The A54S reversion mutation was predicted by PremPS to decrease mAb stability (calculated as ΔΔG) for all four mAbs, indicating the forward mutation (S54A) in the mature FDA-approved sequences was predicted to be stabilizing. In addition, we see on average a 1.9-fold reduction in peptide:MHC-II affinity for peptides from FDA-approved mAbs with this mutation. Thus, *in silico* methods predict that S54A reduces the potential for CD4+ T cell immunogenicity and increases global stability in these four mAbs.

**Figure 7.**
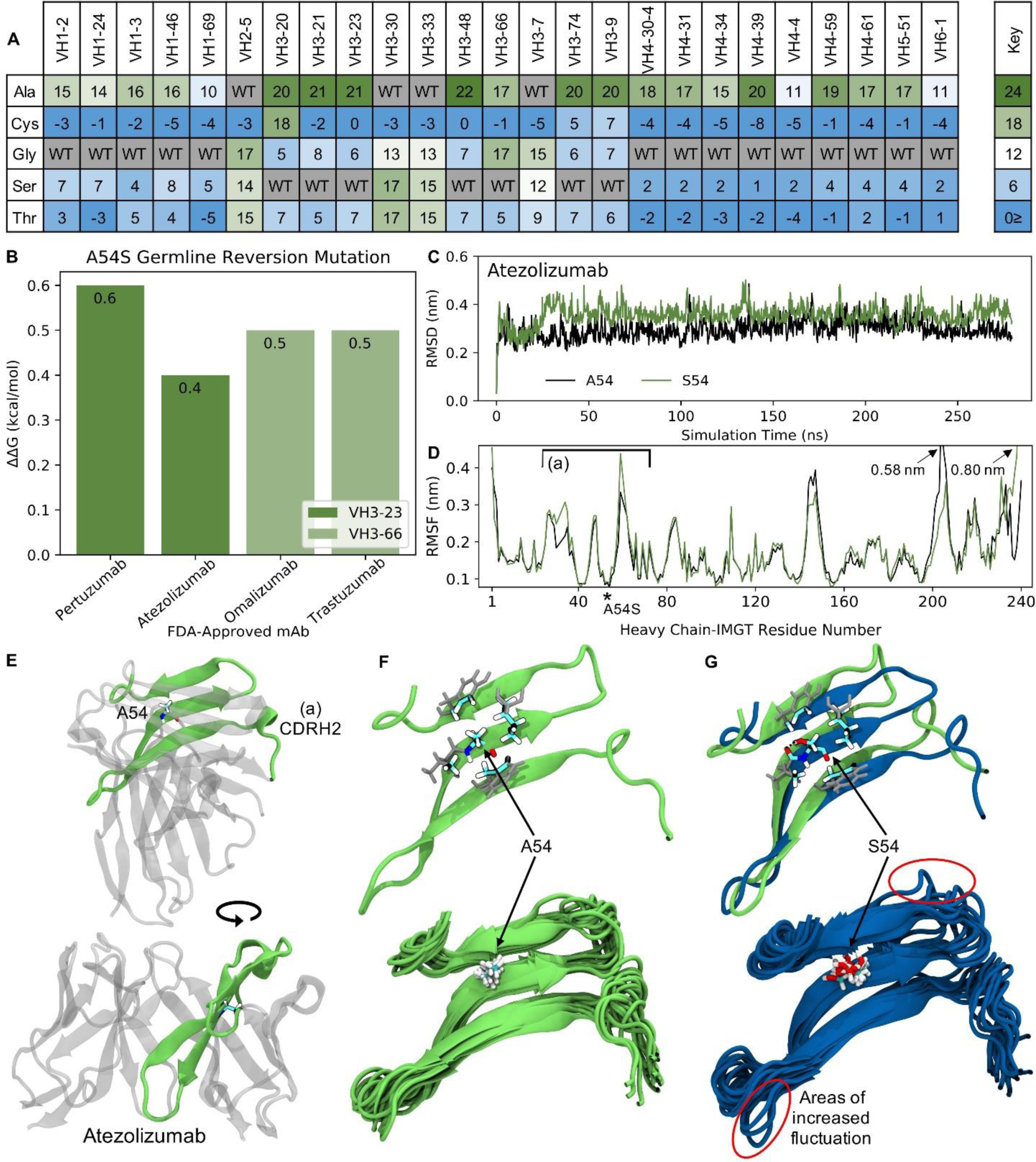
Molecular dynamics (MD) simulations indicate the ‘universal’ FR mutation S54A improves mAb stability. (A) Heatmap of FR scores for the mutation of residue 54 to different amino acids for each of the 25 germline genes. (B) Change in stability as predicted by PremPS upon introducing the A54S germline-reverted mutation into the sequences of four FDA-approved mAbs from two different germlines. (C) Root mean square deviation (RMSD) of Atezolizumab with and without the germline-reverted mutation A54S, referenced to the respective energy-minimized structures, as a function of MD simulation time. (D) Root mean square fluctuation (RMSF) of individual residues in Atezolizumab with and without the A54S mutation, calculated from the MD trajectories. (E) Energy-minimized structure of Atezolizumab in white, with regions in green indicating residues that experienced relatively large fluctuations in the MD simulations, as indicated in (D) by square brackets. (F) Zoomed-in view of region-of-interest (a) from panels (D) and (E), showing the favorable hydrophobic binding pocket of A54 in Atezolizumab (top) and the stability of surrounding loop regions (bottom; 15 overlaid snapshots, equally spaced across the entire trajectory). (G) MD simulation snapshot of Atezolizumab with (blue) and without (green) the germline-reverted mutation A54S, highlighting differences in local secondary structure (top) and relatively large fluctuations in surrounding loops (bottom). A black dashed line indicates a hydrogen bond.

To gain a more mechanistic picture of how the S54A mutation mediates stability in the mature FDA-approved mAbs, we performed molecular dynamics (MD) simulations of Atezolizumab both in its mature form (e.g., with A54) and with the germline-reverted mutation (e.g., with S54). Production simulations were performed for 0.28 *μs* in the NPT ensemble, using the GROMACS-2021 (29,30) simulation engine and the AMBER99SB-ILDN (32) and TIP3P (31) force fields to describe the mAbs and solvent, respectively (see **Methods**). **Figure 7C** shows the root mean square fluctuation (RMSD) of the mature and mutant forms of Atezolizumab from the respective energy-minimized crystal structures as a function of simulation time. The results show the RMSD of the mature mAb was consistently lower than that of the mutant mAb, indicating increased stability of the mature mAb due to the S54A mutation, in line with both our mutation scores and PremPS results.

Increased stability due to the S54A mutation is further supported by the higher root mean square fluctuation (RMSF) of residues in the mutant versus mature mAb during the simulations (**Figure 7D**). In particular, we observe a distinct region in the V_H_ where residues in the mutant mAb experienced increased flexibility (decreased stability) during the simulations, namely IMGT-aligned V_H_ residues 26-72. This region corresponds to a *β*-sheet on which residue 54 is situated (**Figure 7E**), indicating the observed changes in RMSF are directly attributable to this mutation. In the mature mAb simulation, we observe A54 to be optimally situated within a hydrophobic pocket formed by the inward-facing side chains of surrounding residues (**Figure 7F**, top). Favorable binding interactions between the methyl group on A54 with these hydrophobic side chains during the simulation results in relatively small fluctuations of nearby loops (**Figure 7F**, bottom), which includes the CDRH2 loop that is often in direct contact with antigens. Conversely, substituting a serine into this hydrophobic pocket leads to S54 having an unsaturated hydrogen bond. The polar side chain of S54 is observed to rotate freely during the simulation, forming a hydrogen bond only transiently with a nearby residue (**Figure 7G**, top) and causing increased fluctuations in the nearby loops (**Figure 7G**, bottom). The decreased motion of the CDRH2 region due to the S54A mutation is consistent with previous studies showing Ab rigidification may selectively occur during affinity maturation for certain GL families upon binding antigen (51). Future computational studies could test this directly through MD simulations of FDA-approved mAbs in complex with their cognate antigens. **Figure S10** demonstrates this mechanism of stability due to the S54A FR mutation is the same for V_H_3-66-based mAbs as well, underscoring the ‘universal’ nature of this mutation.

We observed a second mutation (Y103F) with even higher prevalence among natural human repertoire antibodies. **Figure 8A** shows the predicted scores for the mutation across residues with similar hydrophobic side chains or ring structures (F, H, M, W and Y) for all of the 25 GLs studied. Our results show that this mutation broadly occurs across many of the GLs, with high scores ranging between 19 and 23.

**Figure 8.**
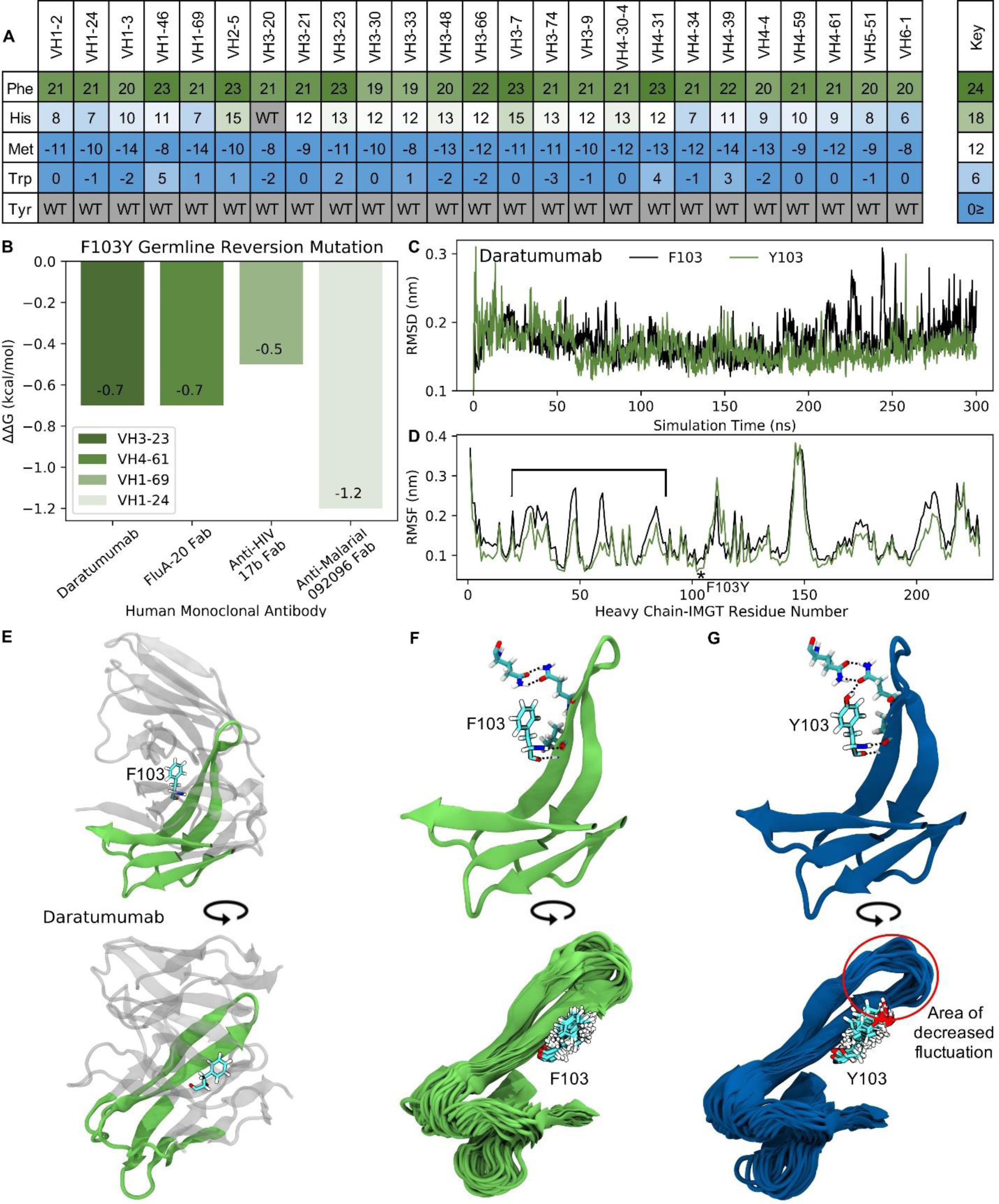
MD simulations indicate the ‘universal’ FR mutation Y103F decreases mAb stability. (A) Heatmap of FR scores for the mutation of residue 103 to different amino acids for each of the 25 germline genes. (B) Change in stability as predicted by PremPS upon introducing the F103Y germline-reverted mutation into the sequence of Daratumumab and three other human mAbs found to have the F103Y mutation. (C) Root mean square deviation (RMSD) of Daratumumab with and without the germline-reverted mutation F103Y as a function of MD simulation time. (D) Root mean square fluctuation (RMSF) of individual residues in Daratumumab with and without the F103Y mutation. (E) Energy-minimized structure of Daratumumab in white with highly fluctuating regions in green, as indicated in (D) by square brackets. (F) Zoomed-in view of region-of-interest from panels (D) and (E), highlighting the reduced hydrogen bonding capabilities of F103 in Daratumumab (top), leading to increased fluctuations in surrounding loop regions (bottom). (G) MD simulation snapshot of Daratumumab with the F103Y mutation, highlighting the strong hydrogen bonding capacity of Y103 (top), leading to reduced fluctuations in surrounding loops (bottom).

To determine the impact of the Y103F mutation on mAb stability, we analyzed the single FDA-approved mAb (Daratumumab) containing this mutation, as well as three other mAbs identified through an IMGT BlastSearch to have this mutation. The three non-FDA-approved mAbs included the anti-influenza FluA-20 Fab from the V_H_4-61 GL family, the anti-HIV 17b Fab from the V_H_1-69 GL family, and the anti-malarial 092096 Fab from the V_H_1-24 GL. As with the S54A mutation, we first utilized PremPS to assess the change in mAb stability upon reversion of the phenylalanine at position 103 in the mature sequences of these mAbs to its identity of a tyrosine in all of the respective germlines. **Figure 8B** shows that this mutation, in contrast with the S54A mutation, is predicted to be destabilizing to all four of the analyzed mAbs. This prompted us to explore this mutation further: why would an antibody repeatably select a destabilizing mutation during affinity maturation, particularly one far away from the binding site?

To address this question, we performed MD simulations of Daratumumab both with and without the Y103F mutation. Production simulations were carried out as described earlier for the simulations with Atezolizumab and Omalizumab (see also **Methods**). Over the converged, second half of the simulation, we observe the RMSD of the mature antibody to be consistently higher than that of the mutant mAb (**Figure 8C**), suggesting this mutation is indeed destabilizing. This result aligns with our PremPS results but still contradicts our mutation scores.

Decreased mAb stability due to the Y103F mutation is also evidenced by the decreased RMSF values of residues in the mutant versus mature mAb, averaged over the converged portion of the simulations (**Figure 8D**). The largest RMSF differences are observed for IMGT-aligned V_H_ residues 19-80. This region corresponds to FR2, CDR2 and FR3 and directly contacts the Y103F mutation, which is situated between the V_H_ and VL domains of the antibody Fab (**Figure 8E**). In the simulation of the mutant mAb, the tyrosine is observed to form consistent hydrogen bonds with neighboring residues, whereas the phenylalanine in the simulation with the mature mAb is unable to do so (**Figures 8F, G,** top). Consistent hydrogen bonds formed by Y103 essentially lock the tyrosine into place, leading to relatively small fluctuations in surrounding loops during the simulation (**Figure 8G**, bottom). Conversely, large fluctuations in neighboring loops are observed in the simulation with the mature mAb (**Figure 8F**, bottom).

Overall, our simulations support the PremPS result that Y103F is a destabilizing mutation. However, due to the mutation’s unique location at the intersection between the V_H_ and VL domains of the antibody, we hypothesize that Y103F may facilitate a favorable conformational transition of the antibody upon antigen binding. This could explain the high prevalence with which this mutation is observed in natural human antibody repertoires and should be explored in future computational studies.

## 1 Discussion

In this work, we generated PSSMs using repertoire sequences from 25 GL genes. We used these scores to assess framework mutations of FDA-approved therapeutic antibodies and to determine the degree of similarity with natural antibody sequences. We found that mutations in FDA-approved antibodies are also common in human antibody repertoires, suggesting these mutations are conferring benefits to the protein.

To understand why high frequency repertoire mutations are beneficial, we analyzed various high-scoring mutations for stability effects, as we expect thermostability to be a top priority to select for in both natural antibodies as well as antibody therapeutics. Due to scant evidence in literature, we were unable to make firm conclusions about this claim. A comprehensive study of stability would be necessary to resolve this issue by directly testing many antibody variants *in vitro*. However, we expect that there should be reasonably high correlation between PSSM scores and stability.

The potential for CD4+ T cell immunogenicity is another critical factor that should be selected for in natural and human-engineered antibodies. Our work shows that increasing PSSM scores correlate with decreasing peptide:MHC-II binding affinity, signifying removal of CD4+ T cell (MHC-II) epitopes and potentially decreasing immunogenicity, a critical quality in antibody therapeutics. In future therapeutic mAb development, repertoire-based PSSMs could be utilized as early warning signals that a sequence may have developability flags or induce an immune response.

The high scoring FR mutation S54A was evaluated *in silico* and by simulations, where we found clear evidence supporting the claim that S54A confers benefits to both stability and reduced MHC-II peptide epitope content.

While we anticipate the factors investigated (thermostability and MHC-II peptide epitope content) to be highly correlated with relative repertoire abundance, there are clearly many other factors to discover. This is made clear by the fact that some mutations that we would expect to be detrimental to stability, like Y103F, seem to have high scores. This must be because they were selected for other benefits not considered in this study. We expect that future research surrounding human antibody repertoires will continue to illuminate the poorly understood selection criteria that are implemented in affinity maturation with regards to framework mutations.

Our work highlights several important findings of natural human antibody repertoires and the dynamics of affinity maturation that produce them. We found that framework mutations are remarkably consistent across individuals, suggesting that affinity maturation is selecting mutations in a consistent manner. This finding indicates that it may be well-suited for computational modeling, allowing for accurate prediction of the types and frequencies of framework mutations produced in affinity maturation. In addition, our results independently confirm the notion that mutation profiles are GL-dependent. We showed that the correlations between mutation scores of germline family members are much higher than those across families. This analysis denotes that mutational frequencies are highly dependent on sequence, likely due to AID’s intrinsic mutational biases. We look forward to future research aimed at determining the degree to which natural human antibody repertoire mutations are dictated by the precursor local sequence environment.

With the current high costs of and increasing demand for mAb therapeutics, methods that can infer sequence to function relationships are very valuable. We hypothesize that incorporating high scoring mutations into mAb sequences will further enhance drug-like properties for existing therapies and could significantly reduce development costs for future mAbs.

## Acknowledgments

The authors wish to acknowledge members of the Whitehead lab for constructive comments.

## Author Contributions

Designed experiments: BMP, SAU, ERR, KGS, MGG, BJD, TAW; Performed experiments: BMP, SAU, MGG; Performed simulations: ERR, KGS; Wrote paper with contributions from all co-authors: BMP, SAU, ERR, KGS, TAW.

## Data Availability Statement

All GL-specific FR scores are given as supplemental datasets. All code used to generate alignments and calculate FR scores are deposited on Github (https://github.com/WhiteheadGroup/FDA_PSSM).

## Contribution to the Field Statement

Monoclonal antibodies are currently the single most important class of biological therapeutics. Due to their increased size and complexity compared to other classes of therapeutics, such antibodies are extremely resource intensive to develop. As such, methods that can infer sequence to function relationships for these therapeutics are very valuable. We analyzed natural human antibody sequences to find high frequency mutations that could enhance properties of antibodies being developed as biological therapeutics. Our findings show that framework mutations (mutations far away from the binding site) in human antibodies are largely repeatable between individuals, suggesting that antibody mutation and selection in humans successfully converges on framework mutations with beneficial characteristics. In addition, we find that antibody therapeutics evolved in a lab setting contain many of these same mutations found in natural human antibodies. We also found that certain development methods produce antibodies that are more “human-like”, and as a result, are less likely to elicit an adverse immune response. We hypothesize that incorporating framework mutations that are observed with high frequency in natural human antibody repertoires into therapeutic antibody sequences will enhance the properties of existing therapeutics and could significantly reduce development costs in the future.

## Supplementary Material

### 2 Supplementary Data

**Data S1.**

PSSMs for each of the 25 analyzed germline genes in csv format.

**Data S2.**

FR scores and number of mutations for 1,000 randomly sampled repertoire antibody sequences across all germlines.

### 3 Supplementary Figures and Tables

#### 3.1 Supplementary Figures

**Figure S1.**
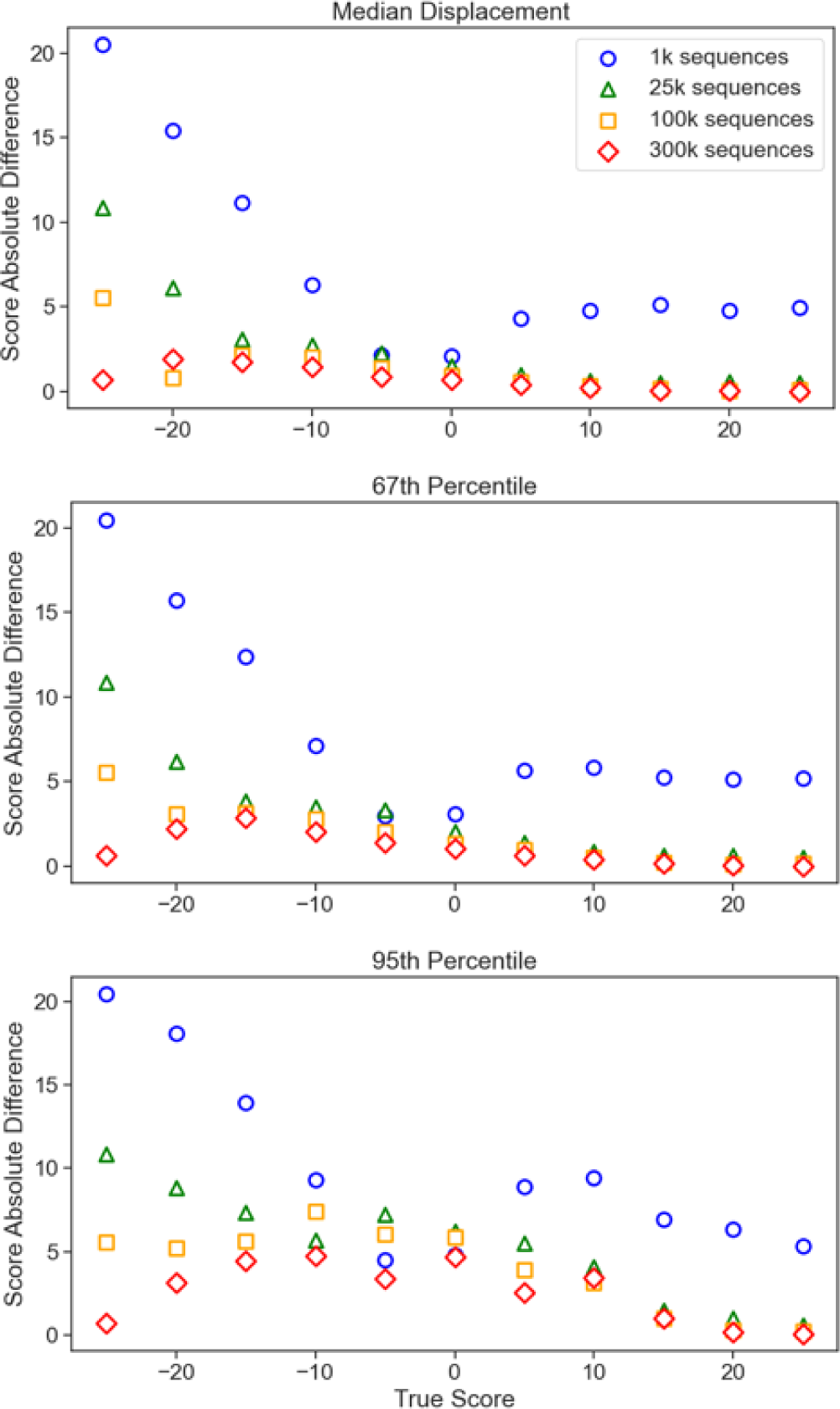
Random subsampling analysis. Simple random samples without replacement of V_H_5-51 repertoire sequences were pulled from the dataset in the above quantities (300,000; 100,000; 25,000; 1,000). Each sample was run through a multiple sequence alignment and a PSSM was generated. Absolute differences between the score using the full number of sequences available and subsample scores were calculated and binned. For each bin, the median, 67^th^ percentile (1σ), and 95^th^ percentile (2σ) values were calculated and plotted. A cutoff was set at 100,000 sequences as the 25,000 and 1,000 sequences resulted in a significant increase in variability, especially at the 95^th^ percentile.

**Figure S2.**
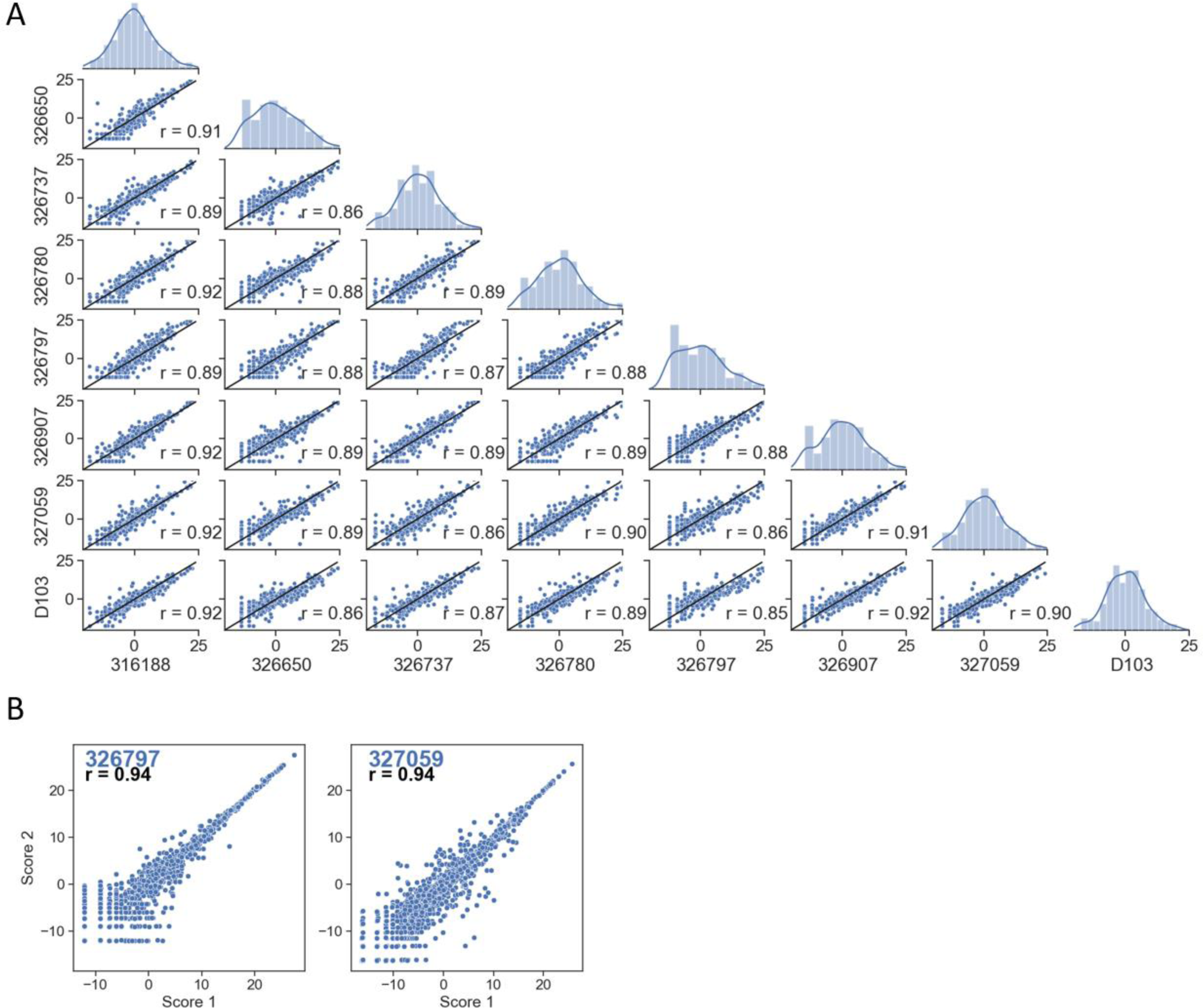
Score correlations for patient-specific PSSMs. (A) Pairwise correlations between scores of patient-specific PSSMs for mutations that are a single nucleotide away from the germline codon. (B) Basal noise for two patients using two independent random samples of 50,000 sequences to generate patient-specific PSSMs. Pearson’s correlation coefficients (r) are calculated for comparisons. Alphanumeric characters on the y- and x-axes represent de-identified subjects.

**Figure S3.**
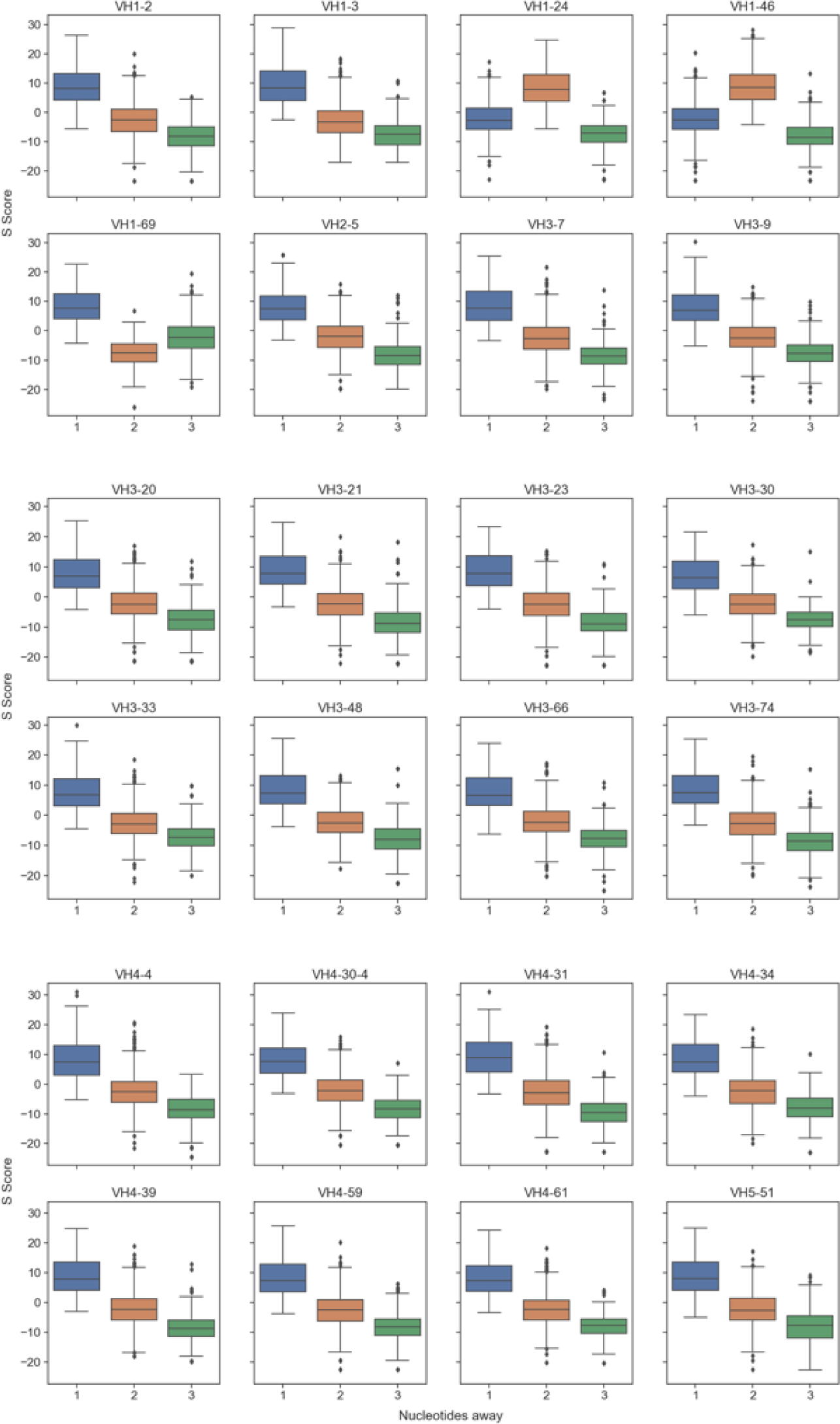
Mutation scores by nucleotide distance from germline. S scores are grouped by minimum number of nucleotide changes from germline to produce the mutation (Blue: 1-nt; Orange: 2-nt; Green: 3- nt). Positions which include allelic variations are not included.

**Figure S4.**
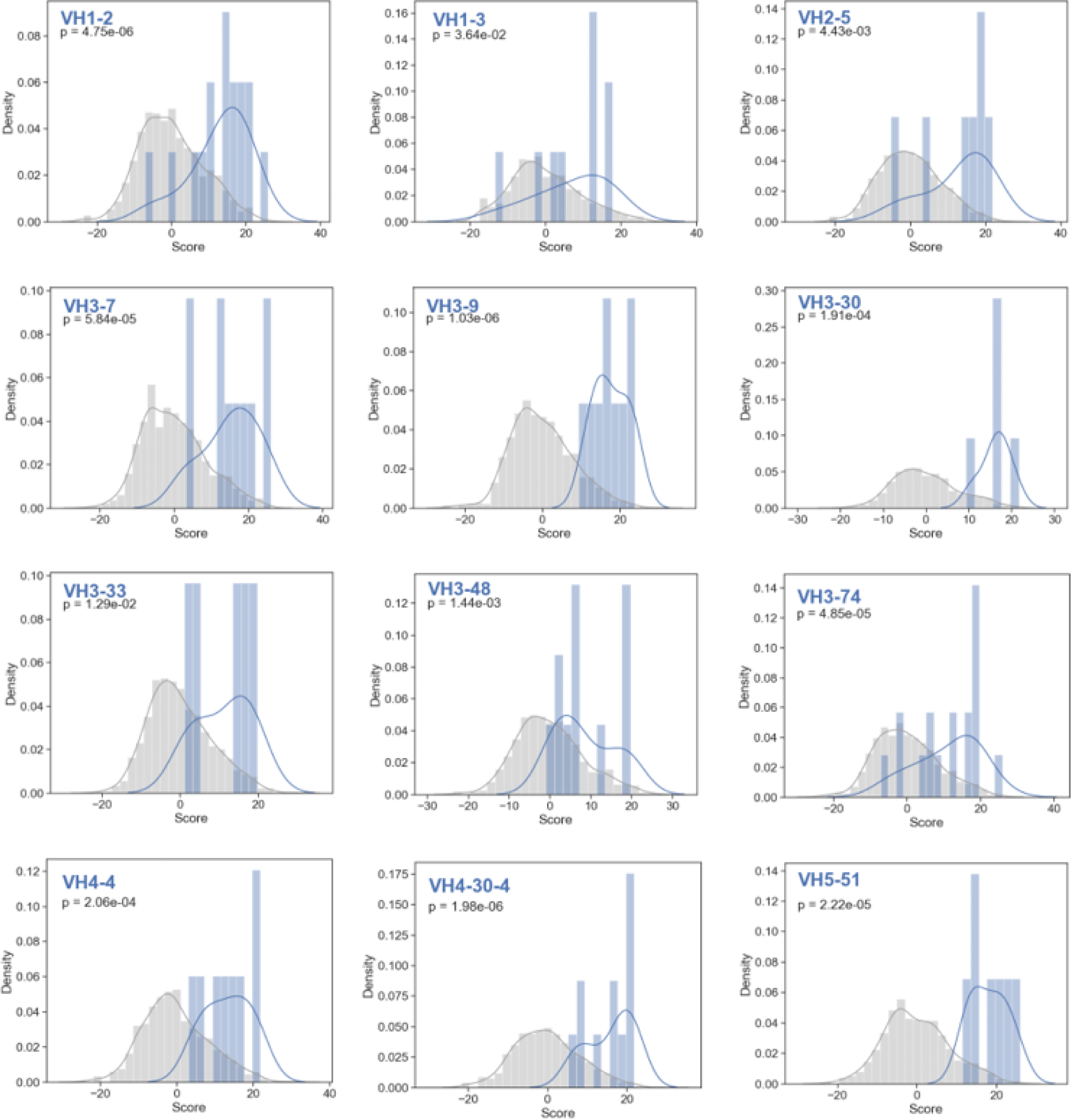
FDA-approved mutation scores compared to their inferred germline gene PSSMs. Histograms of all germline gene PSSMs generated compared to all FDA-approved framework mutations from their respective germlines. P-values are calculated by one-tailed Welch’s t-test.

**Figure S5.**
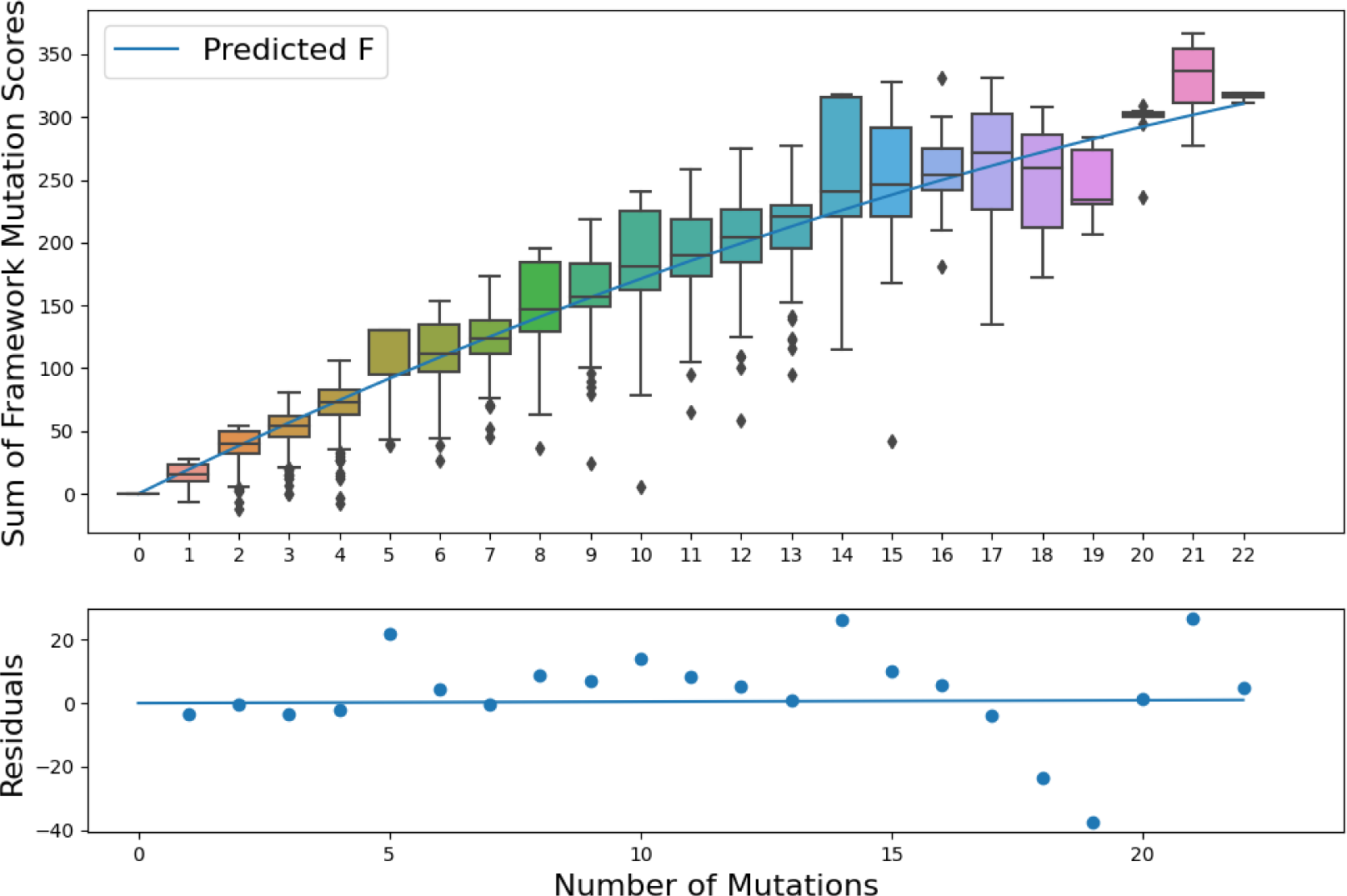
Weighted least squares regression for normalization. Regression plot for an example germline (VH1-46) for a simple random sample without replacement of 10,000 sequences from the repertoire dataset. Average residuals from fitted line are shown below.

**Figure S6.**
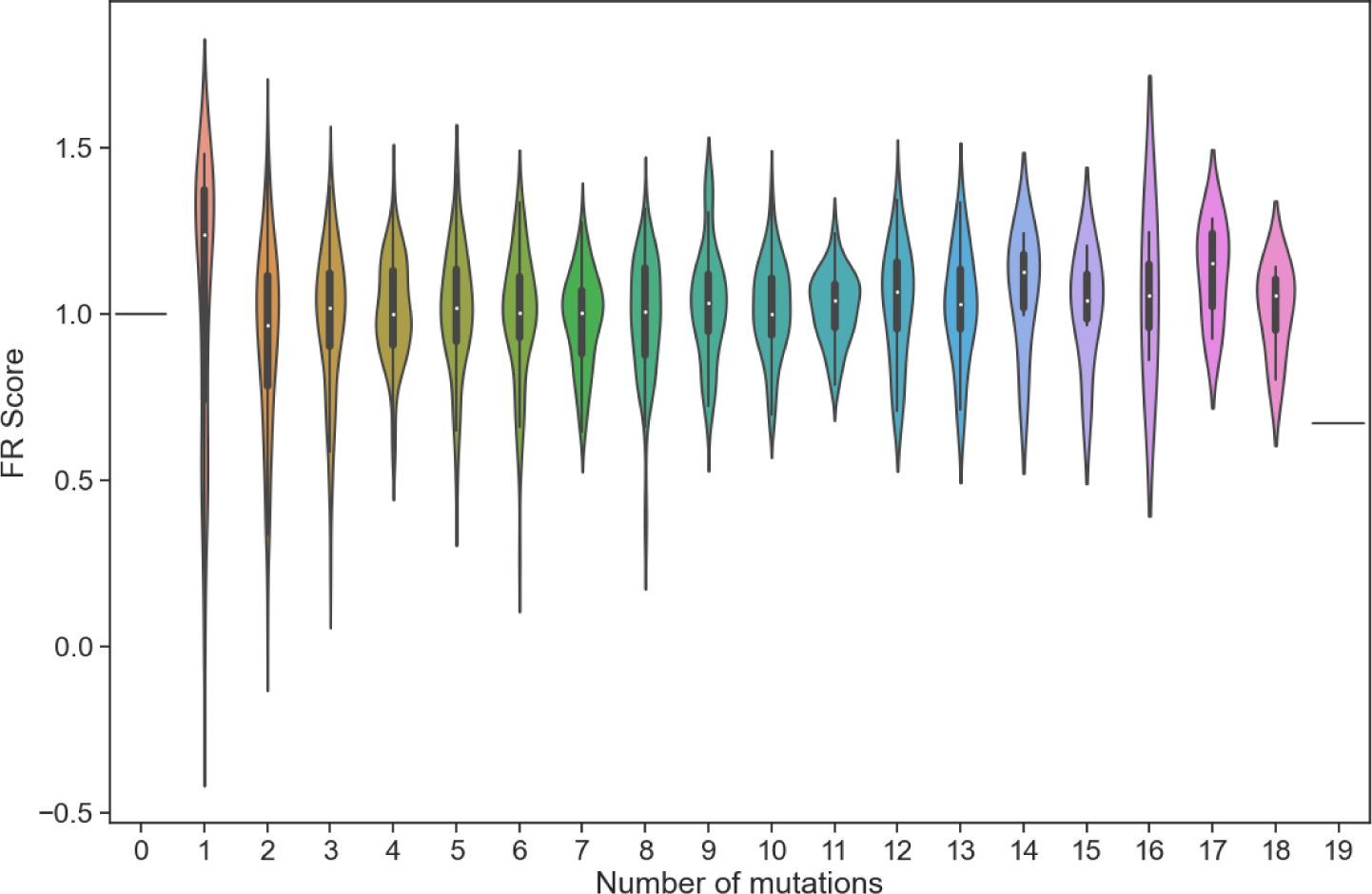
FR score of randomly sampled repertoire antibodies. FR scores for repertoire antibody sequences were calculated from a simple random random sample without replacement of 1,000 sequences across all germlines.

**Figure S7.**
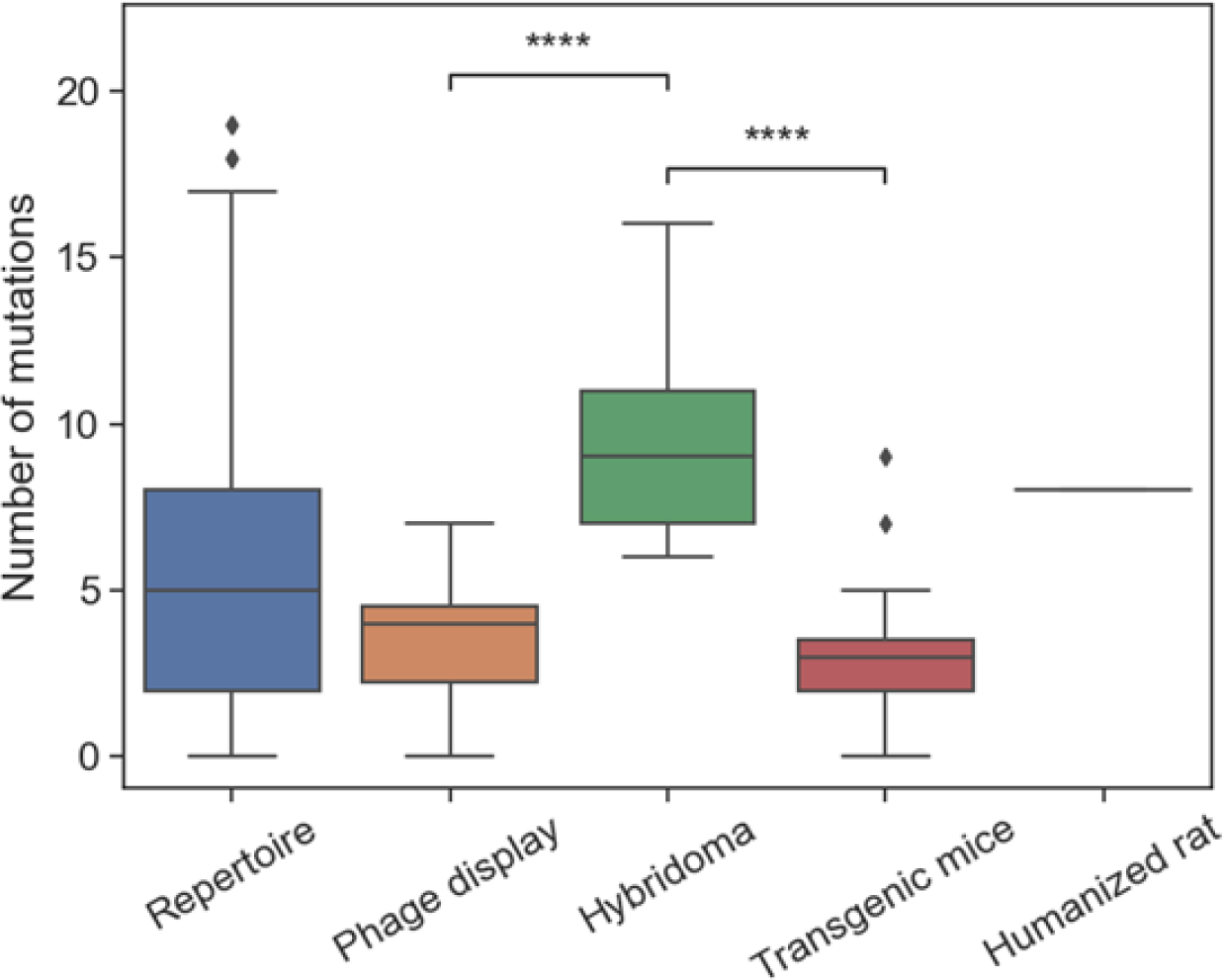
Regulatory approved antibodies developed by hybridomas have a higher number of framework mutations than those developed by other methods. Number of framework mutations for mAbs is shown by development method. (****: p < 1e-4)

**Figure S8.**
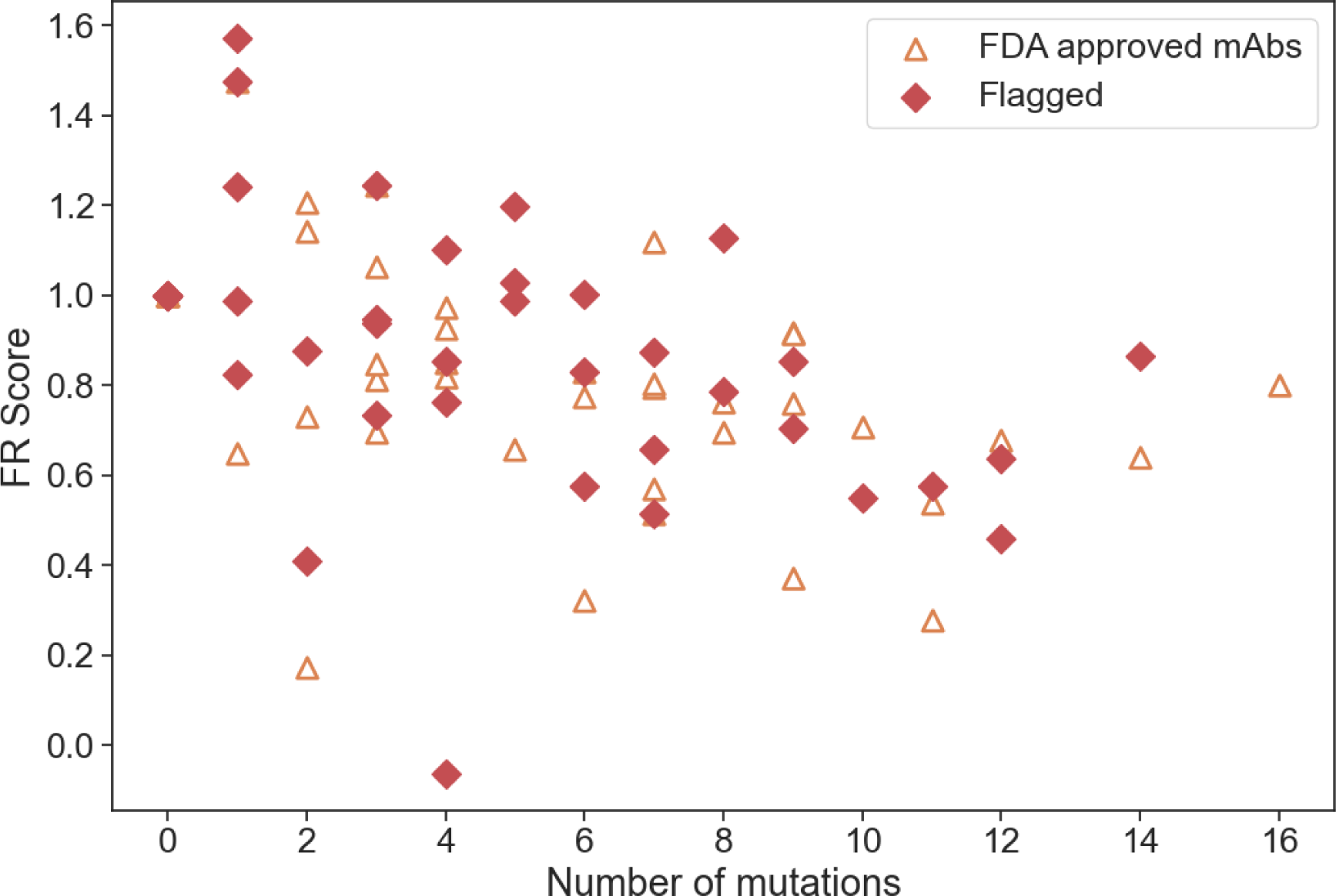
FR scores of FDA-approved and flagged mAbs versus number of framework mutations.

**Figure S9.**
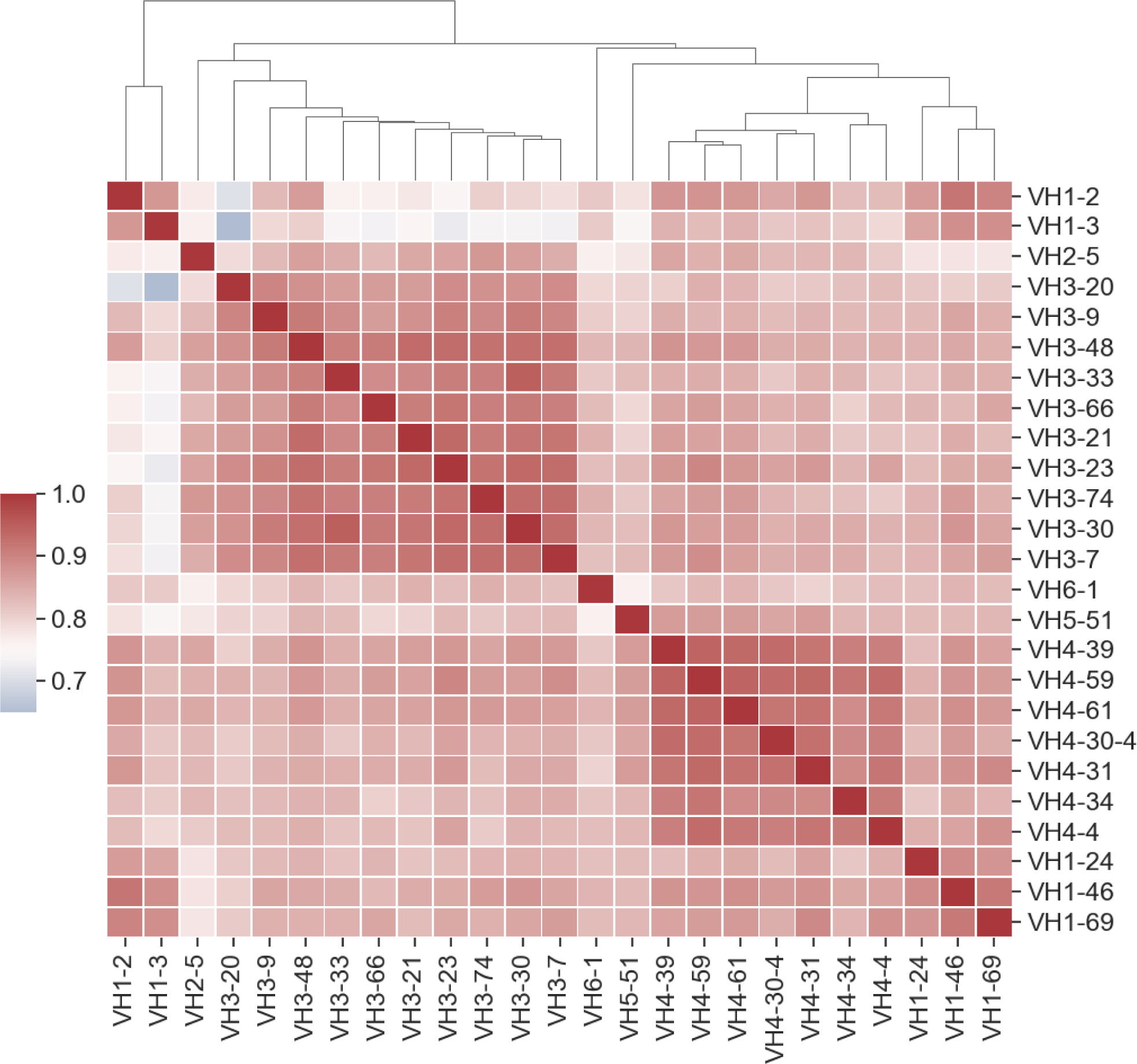
Framework score correlations between germline V_H_ families for shared germline codons only. Heatmap shows correlations between germline V_H_ families where scores are restricted only to shared codons between any two V_H_ germlines. Germline families are grouped by hierarchical clustering of Pearson correlation coefficients. Dendrogram indicates similarity between germline members.

**Figure S10.**
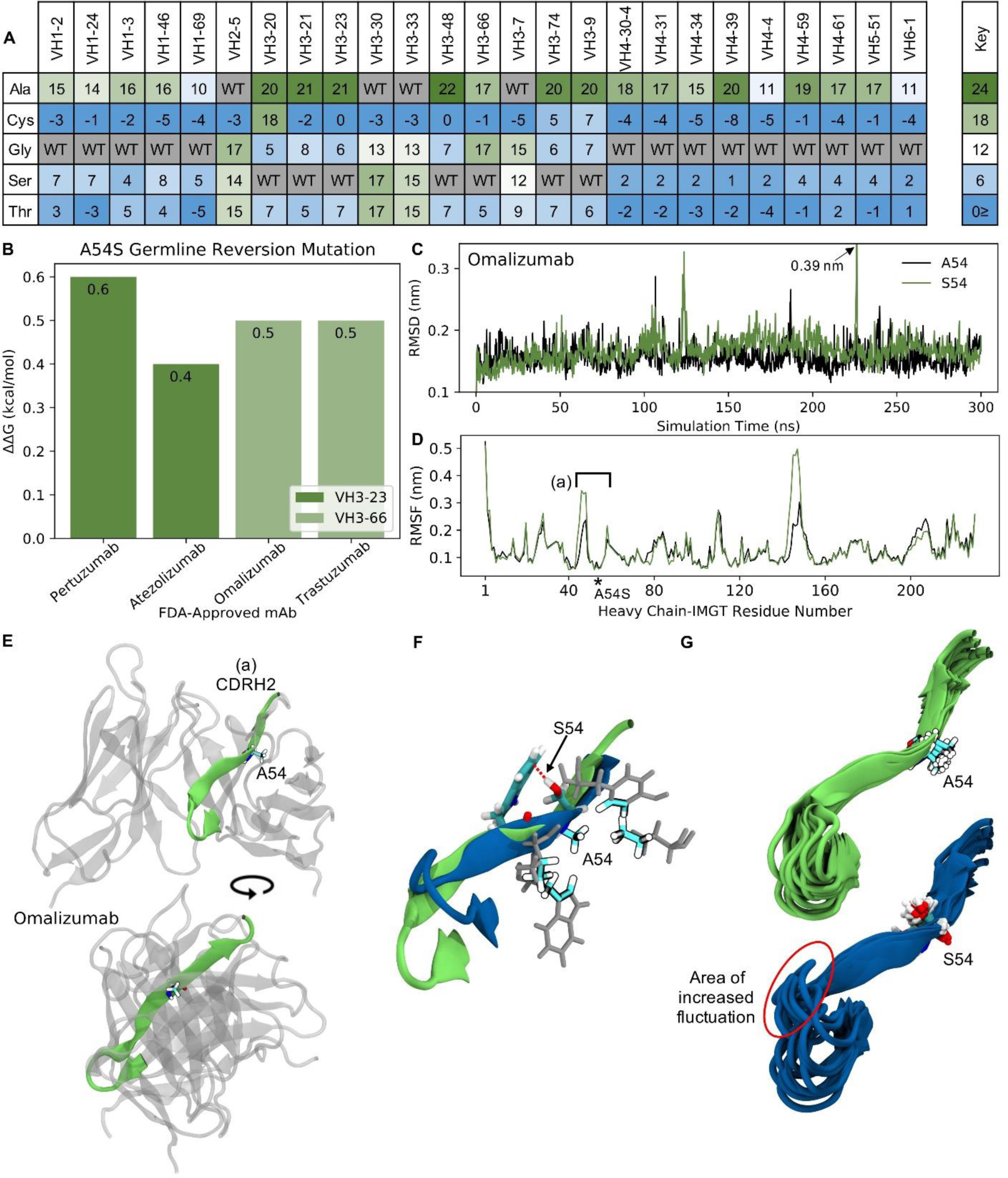
Molecular dynamics (MD) simulations indicate the ‘universal’ FR mutation S54A improves mAb stability in V_H_3-66 germline mAbs. (A) Heatmap of FR scores for the mutation of residue 54 to different amino acids for each of the 25 germline genes. (B) Change in stability as predicted by PremPS upon introducing the A54S germline-reverted mutation into the sequences of four FDA-approved mAbs from two different germlines. (C) Root mean square deviation (RMSD) of Omalizumab with and without the germline-reverted mutation A54S, referenced to the respective energy-minimized structures, as a function of MD simulation time. (D) Root mean square fluctuation (RMSF) of individual residues in Omalizumab with and without the A54S mutation, calculated from the MD trajectories. (E) Energy-minimized structure of Omalizumab in white, with regions in green indicating residues that experienced relatively large fluctuations in the MD simulations, as indicated in (D) by square brackets. (F) Zoomed-in view of region-of-interest (a) from panels (D) and (E), showing the favorable hydrophobic binding pocket of A54 in Omalizumab (secondary structure in green), versus in the mutant simulation (secondary structure in blue), where S54 rotates to the other side of the beta sheet to find a hydrogen bonding partner (red dashed line). (G) Overlaid simulation snapshots (15, equally spaced across the entire trajectory), highlighting increased fluctuations of loops near the S54 mutation in the mutant simulation (bottom) versus in the mature mAb simulation (top).

#### 3.2 Supplementary Tables

**Table S1.** List of all human V_H_ genes and the subset for which a FR PSSM was generated. Exclusion criteria are listed for germlines for which a PSSM was not generated and sequence counts are listed for created PSSMs.

| Germline Gene | PSSM Generated | Exclusion Criteria | Sequences analyzed |
| --- | --- | --- | --- |
| VH1-2 | Yes | N/A | 300,000 |
| VH1-3 | Yes | N/A | 129,913 |
| VH1-8 | No | No associated mAbs | N/A |
| VH1-18 | No | No associated mAbs | N/A |
| VH1-24 | Yes | N/A | 260,282 |
| VH1-38-4 | No | No associated mAbs | N/A |
| VH1-45 | No | Insufficient number of sequences (2,500) | N/A |
| VH1-46 | Yes | N/A | 451,920 |
| VH1-58 | No | No associated mAbs | N/A |
| VH1-68 | No | No associated mAbs | N/A |
| VH1-69 | Yes | N/A | 720,993 |
| VH2-5 | Yes | N/A | 141,372 |
| VH2-26 | No | No associated mAbs | N/A |
| VH2-70 | No | Insufficient number of sequences (24,100) | N/A |
| VH3-7 | Yes | N/A | 2,003,693 |
| VH3-9 | Yes | N/A | 419,523 |
| VH3-11 | No | No associated mAbs | N/A |
| VH3-13 | No | No associated mAbs | N/A |
| VH3-15 | No | No associated mAbs | N/A |
| VH3-16 | No | No associated mAbs | N/A |
| VH3-19 | No | No associated mAbs | N/A |
| VH3-20 | Yes | N/A | 175,018 |
| VH3-21 | No | No framework mutations | N/A |
| VH3-22 | No | No associated mAbs | N/A |
| VH3-23 | Yes | N/A | 300,000 |
| VH3-25 | No | No associated mAbs | N/A |
| VH3-29 | No | No associated mAbs | N/A |
| VH3-30 | Yes | N/A | 1,058,653 |
| VH3-32 | No | No associated mAbs | N/A |
| VH3-33 | Yes | N/A | 710,148 |
| VH3-35 | No | No associated mAbs | N/A |
| VH3-38 | No | No associated mAbs | N/A |
| VH3-41 | No | No associated mAbs | N/A |
| VH3-43 | No | No associated mAbs | N/A |
| VH3-47 | No | No associated mAbs | N/A |
| VH3-48 | Yes | N/A | 300,000 |
| VH3-49 | No | No associated mAbs | N/A |
| VH3-52 | No | No associated mAbs | N/A |
| VH3-53 | No | No associated mAbs | N/A |
| VH3-62 | No | No associated mAbs | N/A |
| VH3-63 | No | No associated mAbs | N/A |
| VH3-64 | No | No associated mAbs | N/A |
| VH3-66 | Yes | N/A | 356,673 |
| VH3-69-1 | No | No associated mAbs | N/A |
| VH3-71 | No | No associated mAbs | N/A |
| VH3-72 | No | No associated mAbs | N/A |
| VH3-73 | No | No associated mAbs | N/A |
| VH3-74 | Yes | N/A | 1,355,540 |
| VH3-NL1 | No | No associated mAbs | N/A |
| VH4-4 | Yes | N/A | 300,000 |
| VH4-28 | No | No associated mAbs | N/A |
| VH4-30-4 | Yes | N/A | 160,757 |
| VH4-31 | Yes | N/A | 373,163 |
| VH4-34 | Yes | N/A | 300,000 |
| VH4-38 | No | No associated mAbs | N/A |
| VH4-39 | Yes | N/A | 300,000 |
| VH4-55 | No | No associated mAbs | N/A |
| VH4-59 | Yes | N/A | 300,000 |
| VH4-61 | Yes | N/A | 280,777 |
| VH5-10-1 | No | No associated mAbs | N/A |
| VH5-51 | Yes | N/A | 702,327 |
| VH6-1 | Yes | N/A | 323,097 |
| VH7-4-1 | No | Insufficient number of sequences (74,500) | N/A |
| VH7-34 | No | No associated mAbs | N/A |
| VH7-81 | No | No associated mAbs | N/A |
| VH8-51-1 | No | No associated mAbs | N/A |

**Table S2.** Allelic variations in V_H_ frameworks for the analyzed germline genes.

| **Germline Gene** | **Allelic Variation(s)** | **Allele** |
| --- | --- | --- |
| VH1-2 | R55W<br>Q69H<br>R75W<br>S78M<br>V100A | VH1-2*02, VH1-2*03, VH1-2*04, VH1-2*07<br>VH1-2*07<br>VH1-2*04<br>VH1-2*02, VH1-2*03, VH1-2*04, VH1-2*05, VH1-2*06, VH1-2*07<br>VH1-2*02, VH1-2*03, VH1-2*04, VH1-2*05, VH1-2*06, VH1-2*07 |
| VH1-3 | K70E<br>T99M | VH1-3*02, VH1-3*03<br>VH1-3*02, VH1-3*03 |
| VH1-46 | F71L | VH1-46*04 |
| VH1-69 | S17A<br>G55R<br>A80T<br>E82K<br>E97D | New allele, VH1-69 germline consensus sequence<br>VH1-69*02, VH1-69*04, VH1-69*07, VH1-69*08, VH1-69*09, VH1-69*11, VH1-69*15, VH1-69*18<br>VH1-69*05, VH1-69*16<br>VH1-69*02, VH1-69*04, VH1-69*06, VH1-69*08, VH1-69*09, VH1-69*10, VH1-69*14, VH1-69*17<br>VH1-69*03 |
| VH2-5 | T11A<br>G40S<br>S68G<br>A100G | VH2-5*08<br>VH2-5*08<br>VH2-5*05, VH2-5*06, VH2-5*09<br>VH2-5*04 |
| VH3-9 | T99M | VH3-9*03 |
| VH3-30 | R17G<br>V55F<br>A68T<br>T77A<br>T86R<br>N92S | VH3-30*02, VH3-30-5*02<br>VH3-30*02, VH3-30-5*02<br>VH3-30*10<br>VH3-30*09<br>VH3-30*13<br>VH3-30*15 |
|  | D98G | VH3-30*05 |
| VH3-33 | H40Y<br>V71A<br>K84T<br>Y88F | VH3-33*07<br>VH3-33*02<br>VH3-33*02<br>VH3-33*02 |
| VH3-48 | A96D | VH3-48*02 |
| VH3-74 | S66T | VH3-74*03 |
| VH4-4 | P16S<br>G17E<br>A24T<br>V25I<br>V42I<br>P46A<br>E55R<br>E55Y<br>I78M<br>K82T<br>C103Y | VH4-4*01, VH4-4*02, VH4-4*07, VH4-4*08, VH4-4*09<br>VH4-4*07, VH4-4*08, VH4-4*09<br>VH4-4*07, VH4-4*08, VH4-4*09<br>VH4-4*04<br>VH4-4*07, VH4-4*08, VH4-4*09<br>VH4-4*07<br>VH4-4*07<br>VH4-4*08, VH4-4*09<br>VH4-4*06, VH4-4*07<br>VH4-4*06, VH4-4*07, VH4-4*08, VH4-4*09<br>VH4-4*02, VH4-4*03, VH4-4*04, VH4-4*05, VH4-4*06, VH4-4*07, VH4-4*08, VH4-4*09 |
| VH4-30-4 | Q17D<br>T24A | VH4-30-4*02<br>VH4-30-4*07 |
| VH5-51 | I39T<br>R43H<br>G47R<br>S83P | VH5-51*02<br>VH5-51*07<br>VH5-51*05<br>VH5-51*04 |

**Table S3.** Normalization constants for FR score. Constants from weighted least squares regression to fit equation of the form: 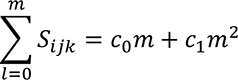 where *m* is the number of framework mutations from germline.

| Gene | c<sub>0</sub> | c<sub>1</sub> | Reduced Chi-Sqr |
| --- | --- | --- | --- |
| VH1-2 | 17.7 | -0.13 | 1.02 |
| VH1-3 | 18.9 | 0.06 | 1.22 |
| VH1-24 | 16.04 | -0.08 | 1.19 |
| VH1-46 | 20.5 | -0.27 | 1.20 |
| VH1-69 | 15.16 | -0.05 | 1.10 |
| VH2-5 | 17.82 | -0.28 | 1.06 |
| VH3-7 | 19.01 | -0.18 | 1.04 |
| VH3-9 | 21.53 | -0.41 | 1.08 |
| VH3-20 | 18.66 | -0.24 | 1.19 |
| VH3-21 | 17.13 | -0.1 | 1.01 |
| VH3-23 | 17.61 | -0.12 | 1.01 |
| VH3-30 | 15.32 | -0.15 | 1.06 |
| VH3-33 | 20.36 | -0.38 | 1.19 |
| VH3-48 | 17.6 | -0.14 | 1.02 |
| VH3-66 | 15.74 | -0.02 | 1.06 |
| VH3-74 | 19.45 | -0.3 | 1.01 |
| VH4-4 | 17.79 | -0.11 | 1.09 |
| VH4-30-4 | 20.63 | -0.43 | 1.34 |
| VH4-31 | 21.72 | -0.32 | 1.39 |
| VH4-34 | 16.48 | -0.08 | 1.01 |
| VH4-39 | 18.01 | -0.2 | 1.03 |
| VH4-59 | 18.06 | -0.18 | 1.01 |
| VH4-61 | 17.05 | -0.21 | 1.04 |
| VH5-51 | 18.36 | -0.21 | 1.13 |
| VH6-1 | 21.1 | -0.49 | 1.54 |

**Table S4.** FR scores and structural coordinates for FDA-approved antibodies. *2 or more nucleotides needed to obtain substitution (from germline and all known alleles)

| FDA Approved Antibody | Germline Gene | Framework Mutations | Mutation Scores | FR Score | PDB ID | Structure Type | Development Method |
| --- | --- | --- | --- | --- | --- | --- | --- |
| Pembrolizumab | VH1-2 | H40Y | 15.0 | 0.84 | 5JXE | Fab complex | Hybridoma |
|  |  | R/W55G | 5.7 |  |  |  |  |
|  |  | Y67F | 18.5 |  |  |  |  |
|  |  | A68N | -6.2* |  |  |  |  |
|  |  | Q/H69E | 17.8 |  |  |  |  |
|  |  | Q72K | 14.9 |  |  |  |  |
|  |  | G74N | 0.0* |  |  |  |  |
|  |  | S/M78L | 20.9 |  |  |  |  |
|  |  | R80T | 16.0 |  |  |  |  |
|  |  | T82S | 10.6 |  |  |  |  |
|  |  | I84T | 17.5 |  |  |  |  |
|  |  | S85T | 21.2 |  |  |  |  |
|  |  | S92K | 9.8* |  |  |  |  |
|  |  | R93S | 25.0 |  |  |  |  |
|  |  | R95Q | 8.4* |  |  |  |  |
|  |  | S96F | 15.3 |  |  |  |  |
| Vedolizumab | VH1-3 | A25G | 17.4 | 0.35 | Unavailable | N/A | Hybridoma |
|  |  | M53I | 12 |  |  |  |  |
|  |  | W55E | -12.1* |  |  |  |  |
|  |  | K66N | 12.8 |  |  |  |  |
|  |  | S68N | -2.9* |  |  |  |  |
|  |  | Q72K | 4.0 |  |  |  |  |
|  |  | I78L | 17.4 |  |  |  |  |
|  |  | R80V | 1.3* |  |  |  |  |
|  |  | T82I | 11.7 |  |  |  |  |
| Benralizumab | VH1-46 | M39I | 28.2 | 0.62 | Unavailable | N/A | Hybridoma |
|  |  | A45R | 1.7* |  |  |  |  |
|  |  | E51A | 8.5 |  |  |  |  |
|  |  | I55Y | -3.3* |  |  |  |  |
|  |  | S66K | 12.2* |  |  |  |  |
|  |  | A68N | -4.5* |  |  |  |  |
|  |  | Q69E | 14.1 |  |  |  |  |
|  |  | K70R | 22.7 |  |  |  |  |
|  |  | Q72K | 12.1 |  |  |  |  |
|  |  | R75K | 5.7 |  |  |  |  |
|  |  | M78I | 14.1 |  |  |  |  |
|  |  | R80S | 18.9 |  |  |  |  |
|  |  | T82R | 9.9 |  |  |  |  |
|  |  | Y103L | 4.4* |  |  |  |  |
| Ravulizumab | VH1-46 | M39I | 28.2 | 0.88 | Unavailable | N/A | Hybridoma |
|  |  | H40Q | 14.1 |  |  |  |  |
|  |  | I55E | 12.8* |  |  |  |  |
|  |  | S66E | 5.0* |  |  |  |  |
|  |  | A68T | 16.8 |  |  |  |  |
|  |  | Q69E | 14.1 |  |  |  |  |
|  |  | K70N | 20.3 |  |  |  |  |
|  |  | Q72K | 12.1 |  |  |  |  |
|  |  | G74D | 20.1 |  |  |  |  |
| Burosumab | VH1-46 | Y67N | 12.6 | 0.62 | Unavailable | N/A | Transgenic mice |
| Risankizumab | VH1-69 | S40H | 12.1* | 0.66 | Unavailable | N/A | Hybridoma |
|  |  | V42M | 8.9 |  |  |  |  |
|  |  | M53I | 12.2 |  |  |  |  |
|  |  | G/R55Y | -9.2* |  |  |  |  |
|  |  | N66K | 18.9 |  |  |  |  |
|  |  | A68N | -2.6* |  |  |  |  |
|  |  | Q69E | 16.8 |  |  |  |  |
|  |  | K70N | 18.9 |  |  |  |  |
|  |  | Q72K | 11.1 |  |  |  |  |
|  |  | R75K | 9.3 |  |  |  |  |
| Ixekizumab | VH1-69 | S40H | 12.1* | 0.77 | 6NOV | Fab | Phage display |
|  |  | G/R55V | 10.1 |  | 6NOU | scFv |  |
|  |  | N66D | 19.2 |  |  |  |  |
|  |  | A68N | -2.6* |  |  |  |  |
|  |  | K70R | 19.1 |  |  |  |  |
|  |  | Q72K | 11.1 |  |  |  |  |
| Galcanezumab | VH1-69 | I39M | 13.7 | 0.53 | Unavailable | N/A | Hybridoma |
|  |  | S40Q | 6.9* |  |  |  |  |
|  |  | G/R55A | 12.4 |  |  |  |  |
|  |  | N66V | 1.7* |  |  |  |  |
|  |  | A68I | 1.7* |  |  |  |  |
|  |  | Q72A | -0.4* |  |  |  |  |
|  |  | G74D | 19.0 |  |  |  |  |
| Palivizumab | VH2-5 | L55D | -3.4* | 0.80 | Unavailable | N/A | Hybridoma |
|  |  | R66D | 4.5* |  |  |  |  |
|  |  | S/G68N | 20.3 |  |  |  |  |
|  |  | T77S | 18.4 |  |  |  |  |
|  |  | T90K | 13.8 |  |  |  |  |
|  |  | M91V | 16.4 |  |  |  |  |
|  |  | V97A | 18.8 |  |  |  |  |
| Secukinumab | VH3-7 | S40N | 24.4 | 0.96 | Unavailable | N/A | Transgenic mice |
|  |  | N55A | 3.7* |  |  |  |  |
|  |  | D69G | 17.3 |  |  |  |  |
|  |  | A96V | 24.9 |  |  |  |  |
| Fremanezumab | VH3-7 | M39I | 13.3 | 0.77 | Unavailable | N/A | Hybridoma |
|  |  | N55E | 3.4* |  |  |  |  |
|  |  | Y66H | 19.3 |  |  |  |  |
|  |  | V68A | 19.9 |  |  |  |  |
|  |  | D69E | 15.2 |  |  |  |  |
|  |  | S70A | 11.5 |  |  |  |  |
| Durvalumab | VH3-7 | None | No Scores | 1 | 5X8M<br>5XJ4 | Fab complex<br>scFv | Transgenic mice |
| Adalimumab | VH3-9 | G55A<br>G66D<br>K72E<br>L101V | 15.6<br>23.7<br>12.5<br>23.3 | 0.94 | 4NYL<br>3WD5 | Fab<br>Fab complex | Phage display |
| Ofatumumab | VH3-9 | G55T<br>N85L | 11.3*<br>19.6* | 0.75 | 3GIZ | Fab | Transgenic mice |
| Sarilumab | VH3-9 | K84E<br>Y88F<br>S93G | 16.0<br>21.4<br>15.4 | 0.87 | Unavailable | N/A | Transgenic mice |
| Ramucirumab | VH3-21 | None | No Scores | 1 | Unavailable | N/A | Phage display |
| Pertuzumab | VH3-23 | S40D<br>S54A<br>A55D<br>Y66I<br>A68N<br>D69Q<br>S70R<br>V71F<br>I78L<br>R80V<br>N82R | 6.7*<br>20.9<br>5.1<br>3.2*<br>-8.2<br>2.2*<br>2.1*<br>-0.1*<br>13.8<br>-0.7*<br>5.6* | 0.28 | 4LLU<br>1S78 | Fab<br>Fab complex | Hybridoma |
| Denosumab | VH3-23 | A55G | 25.6 | 1.46 | Unavailable | N/A | Transgenic mice |
| Daratumumab | VH3-23 | A25V<br>Y103F | 20.6<br>22.6 | 1.24 | 7DUN<br>7DUO | Fab<br>Fab complex | Transgenic mice |
| Avelumab | VH3-23 | S40M<br>A55S<br>Y66F<br>S70T | 4.8*<br>23.4<br>21.2<br>8.4 | 0.84 | 5GRJ | Fab complex | Phage display |
| Dupilumab | VH3-23 | A25G<br>S40T<br>A55S | 17.5<br>23.4<br>23.4 | 1.24 | 6WGB | Fab | Transgenic mice |
| Emicizumab | VH3-23 | M39I<br>S40Q<br>A55S<br>A68R<br>D69R<br>S70E | 14.4<br>1.4*<br>23.4<br>2.7<br>-1.8*<br>-5.4* | 0.34 | Unavailable | N/A | Hybridoma |
| Lanadelumab | VH3-23 | S40M<br>A55G<br>Y66V | 4.8*<br>25.6<br>6.8* | 0.72 | Unavailable | N/A | Phage display |
| Atezolizumab | VH3-23 | M39I<br>S40H<br>S54A<br>A55W<br>R80A<br>N82T<br>L87A | 14.4<br>11.9*<br>20.9<br>-7.7<br>1.8*<br>15.3<br>3.8* | 0.51 | 5X8L | Fab complex | Phage display |
| Ipilimumab | VH3-30 | A54T<br>V101I | 16.7<br>17.4* | 1.14 | 6JC2 | Fab | Transgenic mice |
| Erenumab | VH3-30 | Y67S<br>A68V | 10.9<br>15.7 | 1.05 | 6UMI | Fab | Transgenic mice |
|  |  | Y88F | 20.4 |  |  |  |  |
| Canakinumab | VH3-33 | H40N<br>V55I<br>S93G | 13.6<br>18.6<br>16.3 | 0.84 | 4G5Z | Fab | Transgenic mice |
| Nivolumab | VH3-33 | S22D<br>A24K | 4.7*<br>2.3* | 0.18 | 5GGQ | Fab | Transgenic mice |
| Certolizumab pegol | VH3-48 | V53M<br>S54G<br>Y55W<br>Y66I<br>I78F<br>R80L<br>N82T<br>A83S<br>N85S<br>S86T<br>L87A | 1.5*<br>6.7*<br>5.4*<br>3.6*<br>7.1<br>0.6*<br>12.7<br>17.7<br>19.3<br>18.9<br>1.1* | 0.54 | 5WUV | Fab | Hybridoma |
| Omalizumab | VH3-66 | A25V<br>M39S<br>S40W<br>V42I<br>S54A<br>V55S<br>Y66N<br>N82D<br>L87F | 20.7<br>3.6*<br>-3.7*<br>15.4<br>17.0<br>8.7*<br>12.2<br>16.4<br>14.7* | 0.75 | 4X7S | Fab | Hybridoma |
| Trastuzumab | VH3-66 | M39I<br>S40H<br>S54A | 16.0<br>10.9*<br>17.0 | 0.69 | 6B9Z | Fab | Hybridoma |
|  |  | V55R | 8.0* |  |  |  |  |
|  |  | Y66R | 10.9* |  |  |  |  |
|  |  | R80A | -0.3* |  |  |  |  |
|  |  | N82T | 17.7 |  |  |  |  |
|  |  | L87A | 5.2* |  |  |  |  |
| Eptinezumab | VH3-66 | A25V | 20.7 | 0.90 | Unavailable | N/A | Transgenic mice |
|  |  | S40N | 21.9 |  |  |  |  |
|  |  | S54G | 17.0* |  |  |  |  |
|  |  | D69S | 1.8* |  |  |  |  |
|  |  | S70W | -7.1* |  |  |  |  |
|  |  | V71A | 16.1 |  |  |  |  |
|  |  | N85T | 11.0 |  |  |  |  |
|  |  | L87V | 23.1 |  |  |  |  |
|  |  | Y103F | 22.0 |  |  |  |  |
| Efalizumab | VH3-74 | H40N | 18.8 | 0.66 | 3EO9 | Fab | Transgenic mice |
|  |  | V51E | 18.8 |  |  |  |  |
|  |  | R80V | -2.5* |  |  |  |  |
|  |  | N82K | 5.9 |  |  |  |  |
|  |  | A83S | 17.9 |  |  |  |  |
| Elotuzumab | VH3-74 | H40S | 12.6* | 0.69 | Unavailable | N/A | Hybridoma |
|  |  | V51E | 18.8 |  |  |  |  |
|  |  | V53I | 16.2 |  |  |  |  |
|  |  | S54G | 6.5* |  |  |  |  |
|  |  | R55E | 3.9* |  |  |  |  |
|  |  | S66N | 25.5 |  |  |  |  |
|  |  | D69P | -6.5* |  |  |  |  |
|  |  | V71L | 12.4 |  |  |  |  |
|  |  | G74D | 8.7 |  |  |  |  |
|  |  | R75K | -1.3* |  |  |  |  |
|  |  | T77I | 16.0 |  |  |  |  |
|  |  | T86S | 17.7 |  |  |  |  |
| Tocilizumab | VH4-30-4 | I42V | 18.8 | 0.85 | Unavailable | N/A | Hybridoma |
|  |  | K48R | 17.2 |  |  |  |  |
|  |  | Y66T | 7.7* |  |  |  |  |
|  |  | I78M | 21.1 |  |  |  |  |
|  |  | S79L | 12.1 |  |  |  |  |
|  |  | V80R | 6.5* |  |  |  |  |
|  |  | K90R | 21.3 |  |  |  |  |
| Necitumumab | VH4-30-4 | Y66D | 8.7 | 0.91 | 6B3S | Fab complex | Phage display |
|  |  | I78M | 21.1 |  |  |  |  |
|  |  | L91V | 17.5 |  |  |  |  |
|  |  | S92N | 21.5 |  |  |  |  |
| Alemtuzumab | VH4-4 | G17Q | 4.7 | 0.79 | Unavailable | N/A | Humanized rat |
|  |  | K48R | 16.9 |  |  |  |  |
|  |  | E55F | 6.6 |  |  |  |  |
|  |  | N66E | 9.3* |  |  |  |  |
|  |  | L71V | 15.2 |  |  |  |  |
|  |  | S74G | 20.2 |  |  |  |  |
|  |  | S79L | 12.3 |  |  |  |  |
|  |  | K90R | 21.3 |  |  |  |  |
| Ustekinumab | VH5-51 | I39L | 12.4 | 1.07 | 3HMW | Fab | Transgenic mice |
|  |  | E51D | 14.4 |  |  |  |  |
|  |  | M53I | 13.9 |  |  |  |  |
|  |  | I78M | 18.1 |  |  |  |  |
|  |  | A80V | 25.0 |  |  |  |  |
|  |  | S85T | 21.9 |  |  |  |  |
|  |  | S92N | 20.6 |  |  |  |  |
| Guselkumab | VH5-51 | None | No Scores | 1 | Unavailable | N/A | Phage display |

**Table S5.**
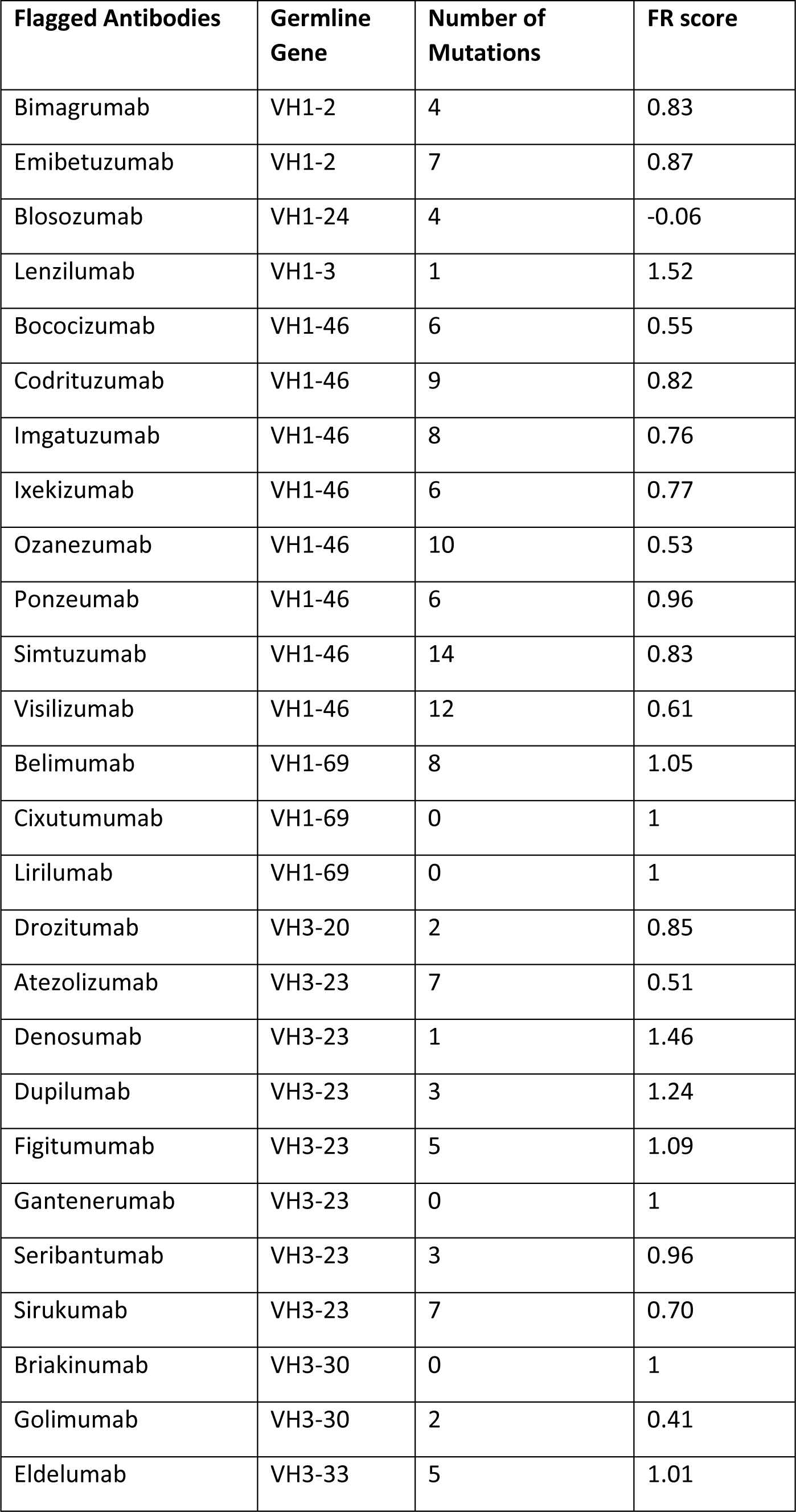
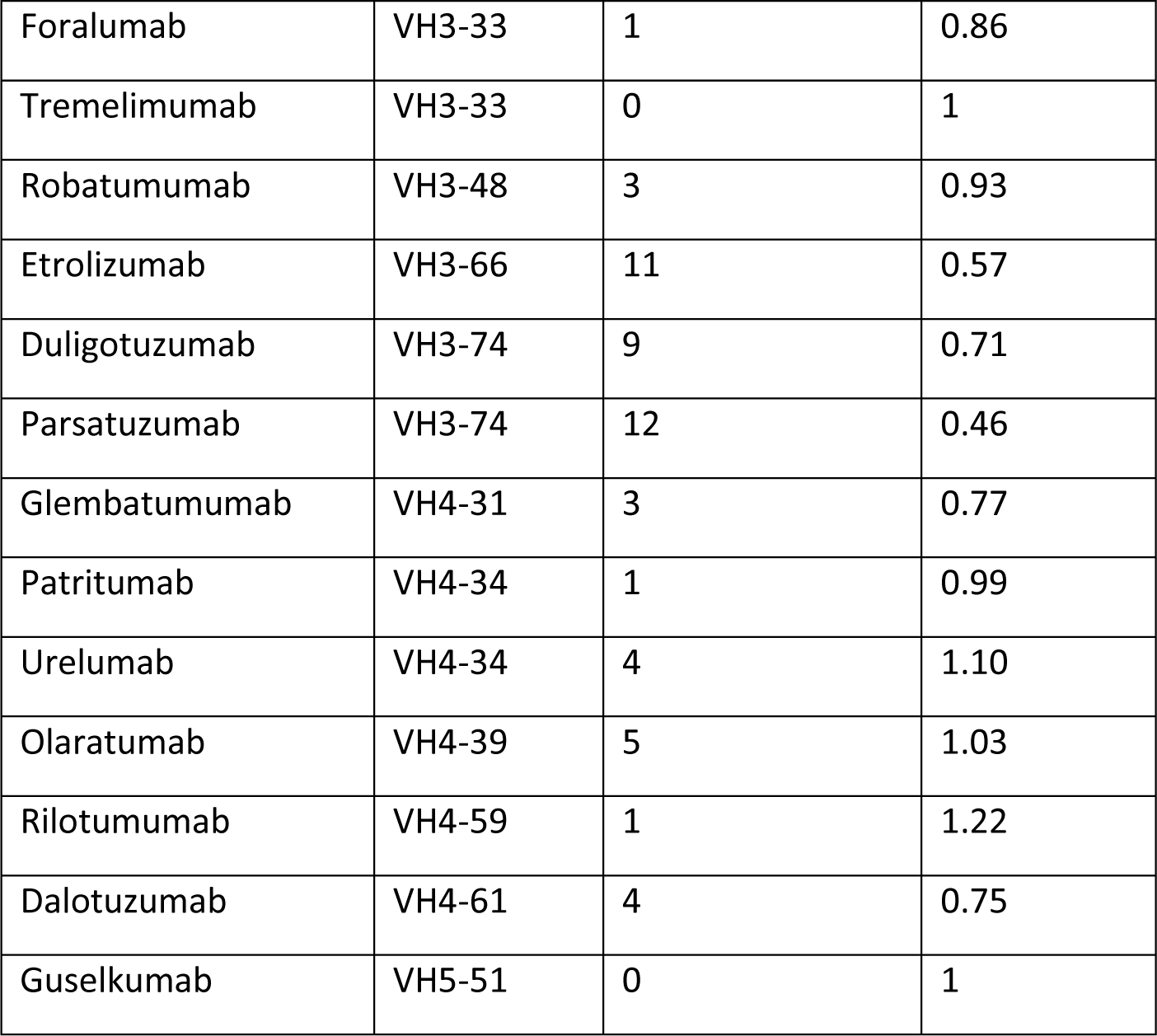
FR scores for flagged antibodies.

**Table S6.**
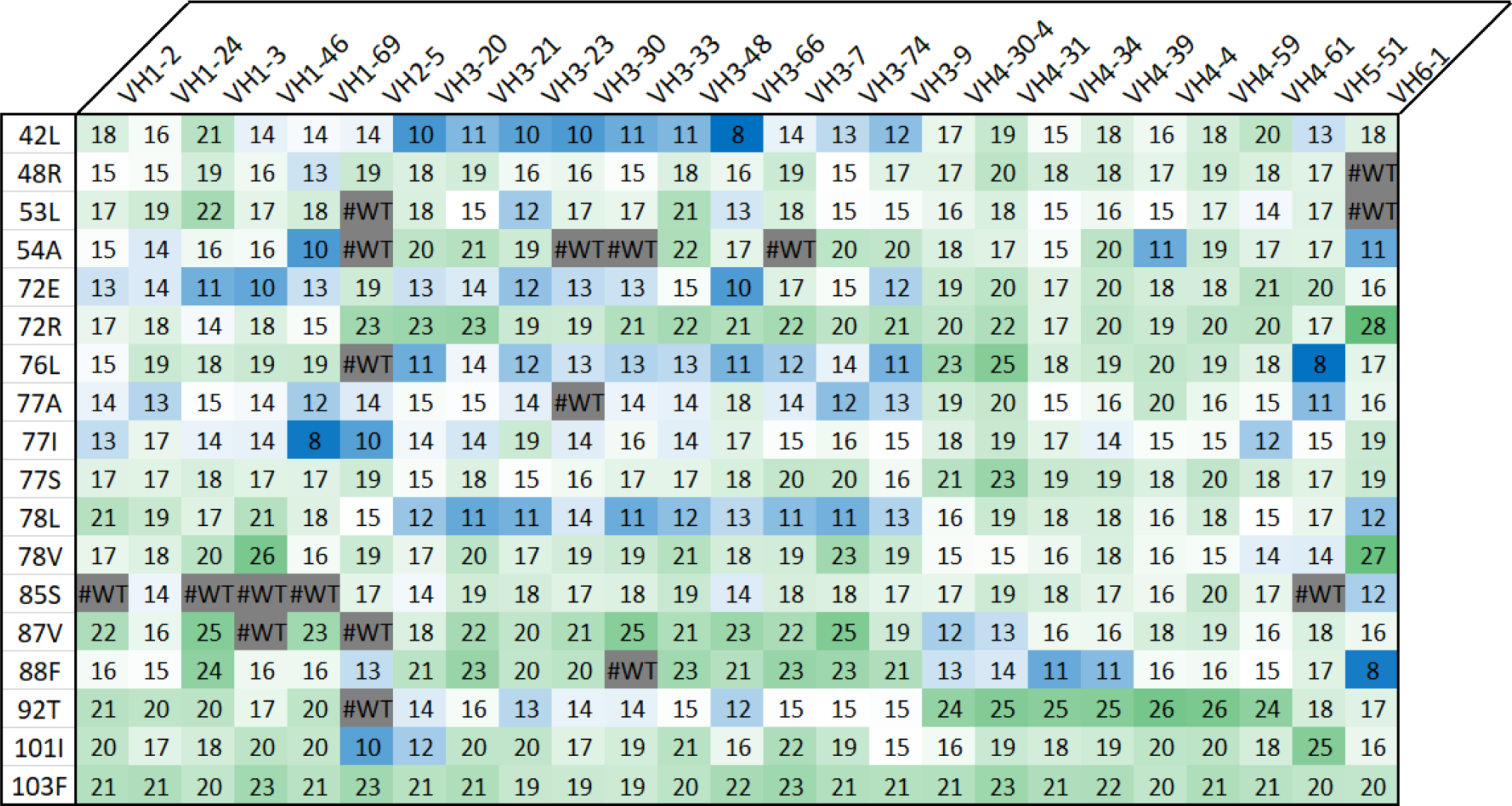
Heatmap of scores across all analyzed germline genes for “universal” FR mutations.

